# PKR inhibition prevents maladaptive responses to endoplasmic reticulum (ER) stress in early-onset dystonia DYT*-PRKRA*

**DOI:** 10.64898/2026.11.09.765397

**Authors:** Emily G. McIntyre, Sravanthi Avula, Taylor Rowson, Julianna C. Aiello, Rekha C. Patel

**Affiliations:** University of South Carolina, Columbia, SC 29208

**Keywords:** Dystonia, PRKRA, PKR, apoptosis, ISR

## Abstract

DYT-*PRKRA* is a rare, childhood-onset dystonia caused by mutations in the *PRKRA* gene, which encodes the protein PACT. PACT activates protein kinase R (PKR) in response to endoplasmic reticulum (ER) stress. In DYT-*PRKRA* patient cells, the PACT-PKR interaction is enhanced, increasing sensitivity to ER stress-induced apoptosis. The canonical protein kinase activated by ER stress is PKR-like endoplasmic reticulum kinase (PERK), which phosphorylates the α subunit of eukaryotic translation initiation factor 2 (eIF2α) to initiate the integrated stress response (ISR) and restore cellular homeostasis via the transcription factor ATF4. However, under persistent stress, ATF4 induces CHOP, shifting the adaptive response toward apoptosis. PKR, also activated by ER stress, phosphorylates eIF2α, but its contribution to cellular homeostasis remains understudied. This work shows, for the first time, that PKR activation is essential for CHOP induction during ISR, directly linking enhanced apoptosis in DYT-*PRKRA* cells to prolonged PKR activation and heightened CHOP induction. Strikingly, a small-molecule PKR inhibitor effectively blocks CHOP expression and significantly decreases apoptosis in DYT-*PRKRA* cells, suggesting PKR inhibition as an effective therapeutic strategy for DYT-*PRKRA*.

**Summary Statement:** This work focuses on understanding the molecular pathomechanism in childhood-onset dystonia DYT-PRKRA, establishing that PKR kinase hyperactivity increases apoptosis in DYT-PRKRA, and suggesting PKR inhibition as a promising therapeutic strategy.

## Introduction

Dystonia is a movement disorder characterized by sustained or intermittent muscle contractions that produce abnormal, repetitive movements and postures (Albanese et al., 2013). More than 35 genes have been identified as causing monogenic inherited dystonia, with new genes and mutations characterized each year (Bragg et al., 2011; Jinnah and Sun, 2019; Gonzalez-Latapi et al., 2021; Thomsen et al., 2024). Among inherited forms of dystonia with childhood onset, consequences are especially severe because abnormal movements emerge during critical periods of motor, speech, cognitive, social, and educational development. DYT-*PRKRA*, previously designated DYT16, is a predominantly pediatric-onset genetic dystonia caused by pathogenic variants in the *PRKRA* gene, which encodes PACT (Camargos et al., 2008; Seibler et al., 2008; Najmabadi et al., 2011; Camargos et al., 2012; Lemmon et al., 2013; Zech et al., 2014; de Carvalho Aguiar et al., 2015; Quadri et al., 2016; Kolbel et al., 2017; Dos Santos et al., 2018; Pinto et al., 2020; Lange et al., 2021; Masnada et al., 2021; Bhowmick et al., 2022; Atasu et al., 2024; Soleymani et al., 2025), the stress-modulated protein activator of protein kinase R (PKR) (Patel and Sen, 1998; Patel et al., 2000). Although adult-onset DYT-*PRKRA* has been reported, most affected individuals develop symptoms in infancy, early childhood, or adolescence (Simon and Patel, 2025). The disorder is frequently progressive and generalized, often producing major disability through impaired gait, painful limb dystonia, speech and swallowing difficulties, and cranial, laryngeal, or oromandibular involvement. Understanding the mechanism of DYT-*PRKRA* is important for identifying molecular targets that could be therapeutically modulated before longstanding, persistent motor disability becomes established.

The original DYT-*PRKRA* cases included children and adolescents with early-onset progressive limb dystonia, followed by generalized dystonia (Camargos et al., 2008; Camargos et al., 2012). Subsequent reports expanded the clinical spectrum to include developmental delay, delayed acquisition of speech or walking, cognitive impairment, bradykinesia, tremor, dysarthria, dysphonia, and, in selected patients, bilateral striatal abnormalities (Simon and Patel, 2025). In several pediatric cases, developmental regression or rapid clinical deterioration occurred after a febrile illness, suggesting that environmental stressors may interact with the underlying *PRKRA* genotype (Lemmon et al., 2013; Kolbel et al., 2017; Pinto et al., 2020; Masnada et al., 2021; Bhowmick et al., 2022). These observations are particularly notable because PACT is a stress-responsive protein involved in the integrated stress response (ISR) (Ito et al., 1999; Patel et al., 2000; Bennett et al., 2004; Bennett et al., 2006; Singh et al., 2009; Singh et al., 2011; Singh and Patel, 2012; Vaughn et al., 2015; Burnett et al., 2020; Frederick and Patel, 2023) and antiviral signaling (Kok et al., 2011; Lui et al., 2017; Chukwurah et al., 2021; Vaughn et al., 2022; Simon and Patel, 2025). Collectively, the clinical phenotype and PACT’s known molecular functions suggest that abnormal responses to cellular stress contribute to disease onset, progression, or both.

PACT functions as a stress-responsive activator of the protein kinase PKR (also known as EIF2AK2) (Clemens and Elia, 1997; Garcia et al., 2006; Dabo and Meurs, 2012; Marchal et al., 2014; Chukwurah et al., 2021) under conditions of cellular stress. PKR is a ubiquitously expressed serine/threonine kinase that stays latent until cellular stress renders it catalytically active. PKR is best known for its role in antiviral defense, where viral double-stranded RNA (dsRNA) activates PKR (Dauber and Wolff, 2009). However, PKR can also be activated independently of dsRNA during nonviral cellular stresses, including oxidative stress, growth-factor deprivation, and endoplasmic reticulum (ER) stress (Ito et al., 1999; Patel et al., 2000; Huang et al., 2002; Bennett et al., 2004; Singh et al., 2009; Singh and Patel, 2012; Farabaugh et al., 2020; Chukwurah et al., 2021). In unstressed cells, PACT is not associated with PKR, and PKR is not enzymatically active. After cellular stress, PACT phosphorylation promotes formation of PACT-PACT and PACT-PKR complexes, activating PKR’s kinase activity. Activated PKR phosphorylates the α subunit of eukaryotic translation initiation factor 2 (eIF2α) at Ser51, thereby activating the integrated stress response (ISR) (Pakos-Zebrucka et al., 2016). The ISR is an evolutionarily conserved pathway that restores protein homeostasis (proteostasis) when cells are stressed (Costa-Mattioli and Walter, 2020). Four distinct stress-responsive kinases are known to phosphorylate eIF2α and initiate the ISR in mammalian cells (Wek et al., 2006): Heme-regulated inhibitor (HRI, also known as EIF2AK1), protein kinase R (PKR, also known as EIF2AK2), PKR-like endoplasmic reticulum (ER) resident kinase (PERK, also known as EIF2AK3), and general control nonderepressible 2 (GCN2, also known as EIF2AK4) (Young and Wek, 2016). Each kinase responds to a distinct set of upstream stress signals with some overlap, yet all converge on the same phosphorylation event, allowing the cell to mount a unified response to diverse forms of cellular stress. HRI is activated by heme deficiency, heavy metal toxicity, heat shock, and osmotic stress (Burwick et al., 2026); PKR is activated during viral infections and other cellular stress signals such as oxidative stress, ER stress, hyperosmolarity, and serum starvation (Patel and Sen, 1998; Patel et al., 2000; Chukwurah et al., 2021); PERK is activated by ER stress (Walter and Ron, 2011); and GCN2 is activated by uncharged transfer RNAs (tRNAs) that accumulate during amino acid deprivation and also by UV irradiation and glucose limitation (Zhu and Marciniak, 2026). The central regulatory event in the ISR is phosphorylation of eIF2α, which transiently attenuates global cap-dependent translation while permitting preferential translation of selected stress-response transcripts, including activating transcription factor 4 (ATF4) (Wortel et al., 2017; Adjibade and Mazroui, 2026). This translational reprogramming reduces protein burden and promotes restoration of cellular homeostasis as ATF4 induces transcription of genes involved in returning cells to homeostasis (Neill and Masson, 2023). The duration and extent of the stress response are regulated, in part, by the level and duration of ATF4 expression. For instance, ATF4 regulates transcription of growth arrest and DNA damage-inducible 34 (GADD34, also known as PPP1R15A), a regulatory subunit of protein phosphatase 1 (PP1) that promotes translational recovery via eIF2α dephosphorylation to promote survival (Kojima et al., 2003; Ma and Hendershot, 2003). Under conditions of continued stress, ATF4 also induces the expression of the transcription factor CCAAT/enhancer binding protein homologous protein (CHOP) (Sano and Reed, 2013). The induction of CHOP and its downstream targets is thought to be a tipping point when the ER stress response becomes pro-apoptotic under chronic stress conditions from which cells cannot recover (Marciniak et al., 2004; Galehdar et al., 2010; Li et al., 2014). Thus, excessive and/or long-term stress results in apoptosis by expression of CHOP and its downstream signaling molecules (Oyadomari et al., 2002). The ultimate cellular outcome depends not only on whether eIF2α is phosphorylated, but also on the timing, amplitude, and resolution of the downstream response. A transient ISR is adaptive; however, delayed, exaggerated, or persistent ISR activation impairs recovery and predisposes cells to apoptosis. This distinction is particularly compelling in a pediatric movement disorder such as DYT-*PRKRA*, in which febrile illnesses and other stressors may coincide with developmental periods of heightened neuronal vulnerability.

Our previous work on the most prevalent DYT-*PRKRA* mutation, P222L, has provided key mechanistic insight into DYT-*PRKRA* (Vaughn et al., 2015). P222L does not substantially impair PACT’s double-stranded RNA binding or abolish its capacity to interact with PKR. Studies in DYT-*PRKRA* patient cells showed enhanced PACT-PKR interaction, increased PACT homodimerization, and abnormal ER stress-dependent PKR activation (Vaughn et al., 2015). In response to ER stress, P222L produced a slower onset but more robust and prolonged PKR activation, accompanied by delayed and persistent eIF2α phosphorylation. These previous observations indicated that P222L is a dysregulating mutation that disrupts the normal temporal control of PACT-mediated PKR activation rather than a simple loss-of-function allele. Compared with control lymphoblasts, P222L patient cells also demonstrated greater apoptosis following ER stress. Therefore, altered activation kinetics of PKR and eIF2α signaling were associated with a functional inability to resolve ER stress appropriately.

In this study, we aimed to further characterize PKR’s contribution to the overall ER stress response and evaluate whether PKR inhibition can protect DYT-*PRKRA* patient cells. PERK is the best-studied kinase activated by ER stress (Korennykh and Walter, 2012; Gardner et al., 2013), and although PKR is also activated by ER stress (Yan et al., 2002; Baltzis et al., 2004; Lee et al., 2007), it has remained unclear how and to what extent PKR contributes to overall cellular survival after ER stress. Because PKR is hyperactive in DYT-*PRKRA* cells and the cells are sensitized to apoptosis, we first characterized both PERK and PKR activation, eIF2a phosphorylation, and ATF4 and CHOP induction in DYT-*PRKRA* patient cells. To understand PKR’s exact contribution to overall cellular homeostasis after ER stress, we used PKR knockout cells and a chemical PKR inhibitor to evaluate ATF4 and CHOP expression. Our results show that PKR activation is crucial for timely, robust eIF2α phosphorylation; ATF4 induction; and ATF4-dependent downstream transcriptional programs after ER stress. In particular, we show that PKR activity is required to induce the pro-apoptotic transcription factor CHOP, linking heightened apoptosis in DYT-*PRKRA* cells to prolonged PKR activation. Importantly, a small-molecule PKR inhibitor blocked CHOP expression and significantly reduced ER stress-induced apoptosis in cells from DYT-*PRKRA* patients. This work advances our mechanistic understanding of regulatory events downstream of PKR activation after ER stress and offers potential future therapeutic strategies for DYT-*PRKRA*.

## Results

We previously reported that compared with wt (normal) lymphoblasts, the homozygous P222L DYT-*PRKRA* lymphoblasts are more susceptible to apoptosis in response to ER stress (Vaughn et al., 2015). In that study, we primarily focused on PACT-mediated PKR activation in DYT-*PRKRA* patient cells after ER stress and did not investigate PERK activation or the molecular events downstream of PKR activation and eIF2α phosphorylation. The current study examines these events to clarify the role of PKR activation in the overall ER stress response and cell survival. Because PERK is the first eIF2α kinase to sense ER stress, we examined whether PERK activation, as measured by the presence of phosphorylated PERK (p-PERK) after treatment with the ER stressor tunicamycin, was different in wt and DYT-*PRKRA* lymphoblasts. To assess PERK activation, we used a well-established method to detect reduced mobility of phosphorylated, catalytically active p-PERK compared with its unphosphorylated, inactive form. As seen in Fig. 1A, PERK was activated equally and with the same kinetics in both wt and DYT-*PRKRA* lymphoblasts (lanes 1-10). We also compared the time course of PKR activation as measured by levels of phosphorylated PKR (p-PKR) in wt and DYT-*PRKRA* patient lymphoblasts under the same conditions. To detect p-PKR, we used a phospho-specific antibody that recognizes PKR phosphorylated at threonine 446, a characteristic of catalytically active PKR. In wt lymphoblasts (left), PKR activation was observed 2 h after tunicamycin treatment, was sustained until 8 h (lanes 2–4) and decreased at 12 h (lane 5). In contrast, DYT-*PRKRA* lymphoblasts (right) showed a much slower increase in PKR activation (lanes 7-9) and, more strikingly, p-PKR levels remained elevated at 12 h after tunicamycin treatment (lane 10). A comparison of eIF2α phosphorylation also showed differences. In wt lymphoblasts (left), we observed a low basal level of eIF2α phosphorylation in the untreated cells (lane 1), followed by increased eIF2α phosphorylation at 2 h that peaked at 4 h (lanes 2-3), declined at 8 h (lane 4), and returned to basal levels by 12 h (lane 5). In contrast, in the DYT-*PRKRA* lymphoblasts (right), we observed a similar increase in eIF2α phosphorylation at 2 and 4 h after treatment (lanes 7-8); however, eIF2α phosphorylation was sustained even at 8 and 12 h post-treatment (lanes 9–10). We quantified p-PKR and p-eIF2α band intensities, normalized them to total PKR and eIF2α, and plotted the ratios in Fig. 1B. The most striking difference is that, unlike in wt cells, both PKR and eIF2α phosphorylation did not return to basal levels in DYT-*PRKRA* cells at 12 h after ER stress, indicating a sustained response to ER stress. Because we observed differences in eIF2α and PKR phosphorylation between wt and DYT-*PRKRA* lymphoblasts, we examined whether the downstream effects of eIF2α phosphorylation also showed differences (Fig. 1C). In wt lymphoblasts (left), ATF4 was undetectable in untreated cells (lane 1), and its expression increased in a time-dependent manner from 2 to 8 h post-treatment (lanes 2–4) and declined at 12 h after treatment (lane 5). In contrast, in the DYT-*PRKRA* patient lymphoblasts (right), although we observed increased expression of ATF4 from 2 to 8 h post-treatment (lanes 7–9), reaching approximately 2.5-fold the levels seen in wt cells, it persisted at high levels even at 12 h (lane 10) post-treatment. Finally, we examined levels of CHOP, an ATF4-induced pro-apoptotic protein, in wt and DYT-*PRKRA* lymphoblasts. CHOP was undetectable in untreated wt cells (lane 1), and its expression increased at 4 and 8 h post-treatment (lanes 3–4) and declined at 12 h after treatment (lane 5). In contrast, in DYT-*PRKRA* patient lymphoblasts (right), we observed markedly higher CHOP expression at 4 and 8 h post-treatment (lanes 8–9), and it continued to rise at 12 h post-treatment (lane 10). Fig. 1D shows quantified band intensities, normalized to the loading control β-actin. Collectively, these results demonstrate dysregulation of the ISR pathway in DYT-PRKRA cells, in which prolonged PKR activation and eIF2α phosphorylation are accompanied by elevated, persistent ATF4 levels and sustained CHOP induction, despite unchanged PERK activation.

**Figure 1.**
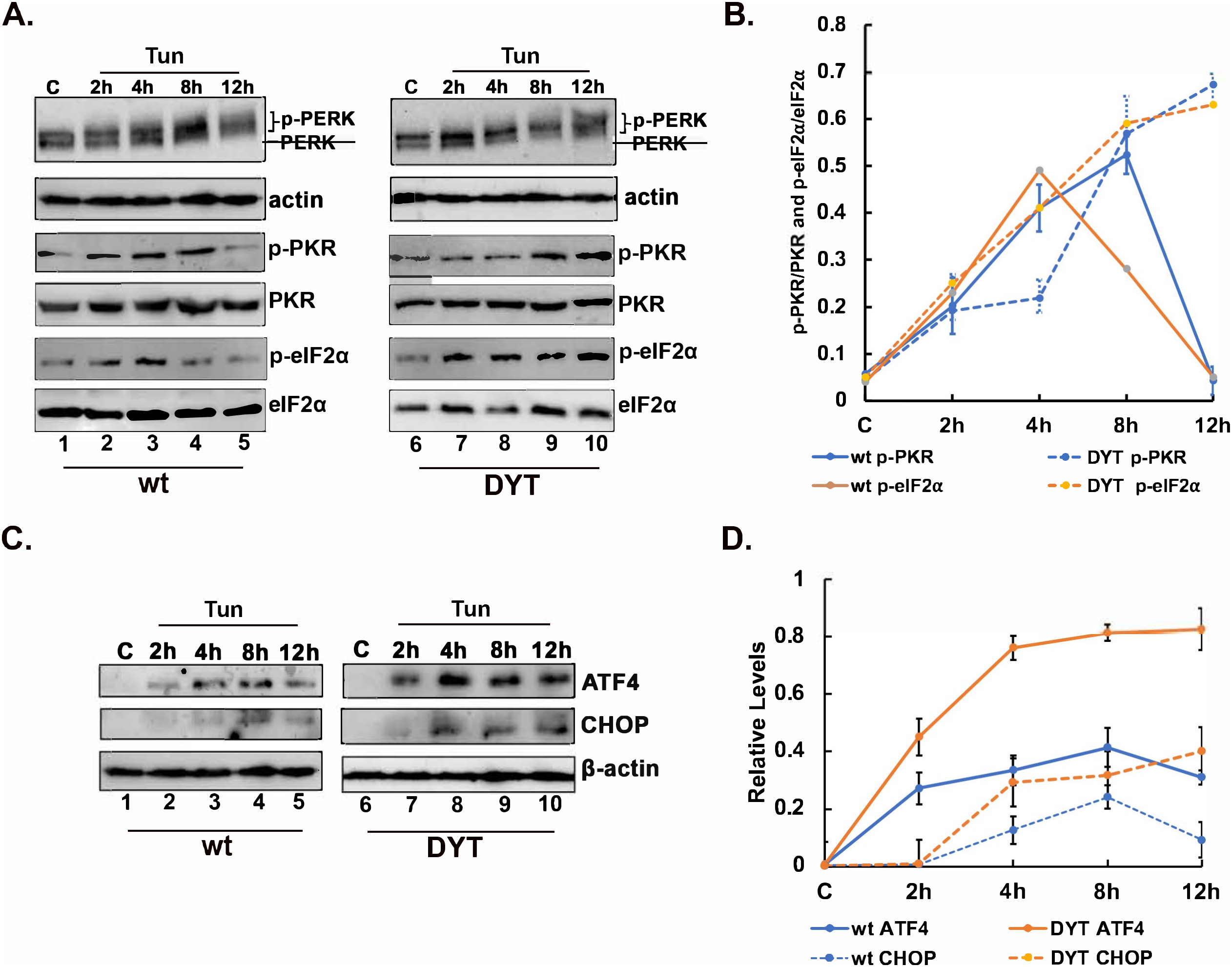
ISR in response to tunicamycin in normal (wt) and DYT-*PRKRA* (DYT) patient lymphoblasts. (A) Western blot analysis for PERK, p-PKR and p-eIF2α. Whole-cell extracts from normal (wt) and DYT-*PRKRA* (DYT) patient-derived lymphoblasts treated with 0.5 μg/mL of tunicamycin (Tun) were analyzed at the indicated time points. Blots were probed for PERK, p-PKR, total PKR, p-eIF2α, and total eIF2α. The best of three representative blots is shown. β-actin was used as a loading control to ensure equal amounts of protein were loaded in each lane. **(B)** The signal intensities of p-PKR, total PKR, p-eIF2α and total eIF2α bands were quantified using ImageQuant TL (Cytiva) and the ratios of p-PKR/PKR and p-eIF2α/eIF2α were calculated for each time point using three separate experiments. The error bars represent standard errors. Solid blue and orange lines: wt; dashed blue and orange lines: DYT-*PRKRA*. Blue lines: p-PKR/PKR and orange lines: p-eIF2α/eIF2α. **(C)** Western blot analysis for ATF4 and CHOP. Whole-cell extracts from normal (wt) and DYT-*PRKRA* (DYT) patient-derived lymphoblasts treated the same as in 1A were analyzed at the indicated time points. Blots were probed for ATF4 and CHOP. The best of three representative blots is shown. We used β-actin as a loading control to ensure equal protein loading in each lane. **(D)** The signal intensities of ATF4, CHOP, and β-actin bands were quantified using ImageQuant TL (Cytiva), and ATF4 and CHOP intensities were normalized to β-actin for each time point using three separate experiments. The error bars represent standard errors. Solid blue: wt ATF4; solid orange: DYT ATF4; dashed blue: wt CHOP; dashed orange: DYT CHOP.

Because Fig. 1 indicates that sustained PKR activation correlates with sustained induction of ATF4 and the pro-apoptotic transcription factor CHOP, we further assessed PKR’s contribution to the ER stress response. We reasoned that if higher and prolonged PKR activation drives higher ATF4 and CHOP expression in DYT-*PRKRA*, then the absence of PKR activation after ER stress would have the opposite effect, driving down both CHOP expression and apoptosis. Thus, we compared the response of PKR+/+ and PKR−/− MEFs to tunicamycin treatment. As shown in Fig. 2A, both PKR+/+ and PKR−/− MEFs showed similar levels and time courses of PERK activation in response to ER stress (lanes 2-5 and 7-10), indicating that PKR absence does not affect PERK activation after ER stress. Similar to wt lymphoblasts, PKR activation was observed at 2-8 h after ER stress in PKR+/+ MEFs (lanes 2-4), and it was reduced at 12 h post-treatment (lane 5). As expected, PKR expression and activation were absent in PKR−/− MEFs (lanes 6-10). The eIF2α phosphorylation in PKR+/+ MEFs followed a pattern similar to that in wt lymphoblasts, with p-eIF2α levels rising at 2 h, peaking at 4 h after ER stress (Fig. 2B, lanes 2-3) and falling at 8 and 12 h post-treatment (lanes 4-5). In contrast, in PKR−/− MEFs, p-eIF2α levels rose at 2 h after ER stress (lane 7), owing to PERK activation, and fell sharply at 4 h, remaining low through 12 h post-treatment (lanes 8-10). These results indicate that in the absence of PKR, although initial eIF2α phosphorylation occurs rapidly after tunicamycin exposure via activated PERK, it is not sustained as in PKR+/+ MEFs, as p-eIF2α levels at 4 and 8 h post-treatment were much lower than in PKR+/+ MEFs. We quantified the p-eIF2αand total eIF2α band intensities in PKR+/+ and PKR−/− MEFs, calculated the p-eIF2α/eIF2α ratios, and plotted them as a bar graph in Fig. 2C. ATF4 expression was strongly induced at 2–12 h after ER stress in PKR+/+ MEFs (Fig. 2B, lanes 2-5). Correspondingly, CHOP expression was seen at 4–12 h after ER stress in PKR+/+ MEFs (lanes 3–5). In contrast, in PKR−/− MEFs, ATF4 induction remained low at 2 and 4 h after ER stress (lanes 7–8), became detectable at 8 h, and increased further at 12 h (lanes 9–10). CHOP was undetectable until 12 h after ER stress, when it appeared weakly (lane 10), at markedly lower than in PKR+/+ MEFs. These results indicate that PKR activation is required for a robust, timely ISR, and that PKR plays a critical role in the rapid induction of ATF4 and its downstream target CHOP. The excessive and prolonged PKR activation seen in DYT-*PRKRA* cells (Fig. 1) is thus likely to directly cause excessive induction of both ATF4 and CHOP.

**Figure 2.**
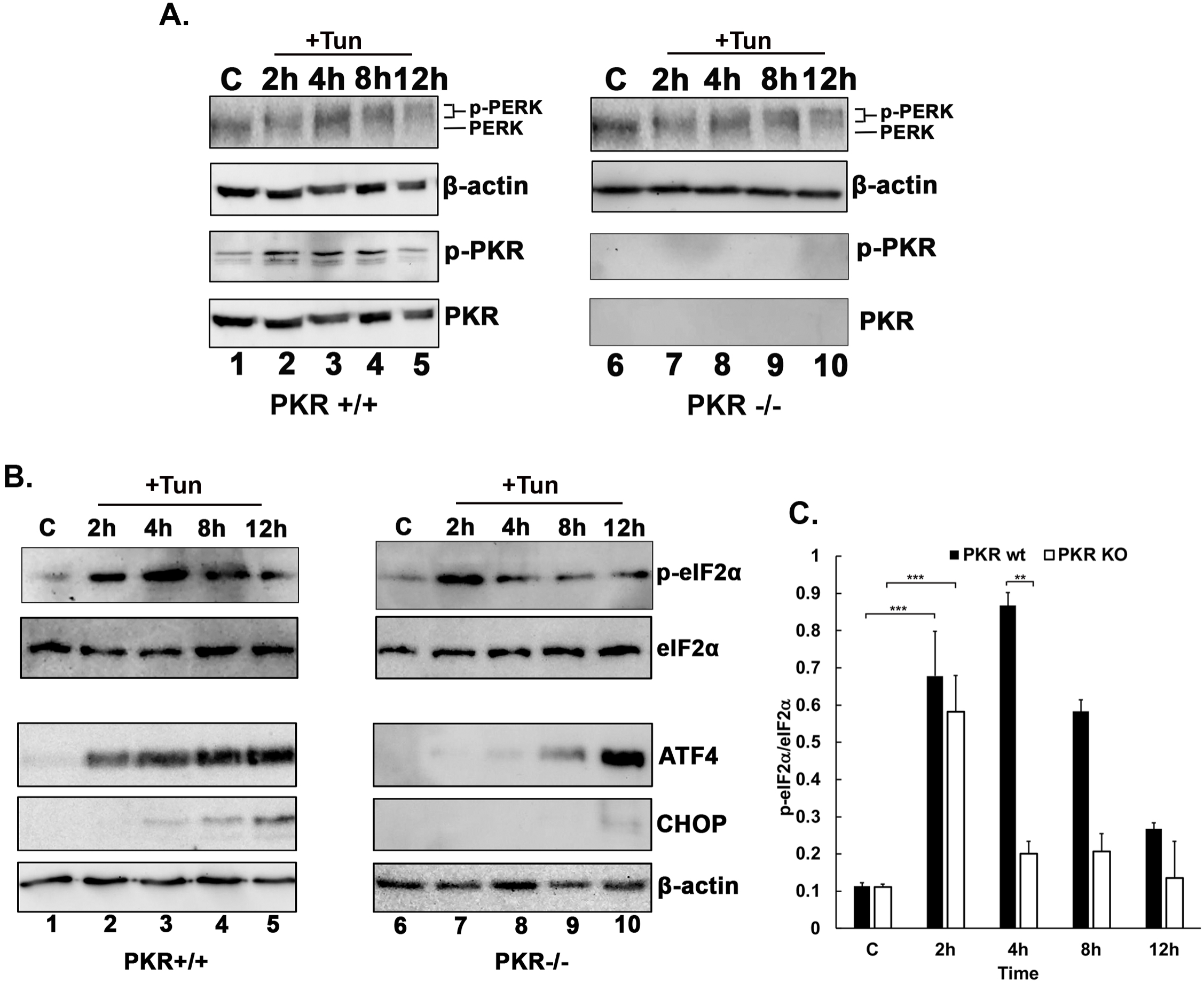
PKR contributes to a timely and robust ISR. (A) Western blot analysis for PERK and p-PKR in PKR+/+ and PKR−/− MEFs. PKR+/+ and PKR−/− MEFs were treated with 0.5 µg/mL tunicamycin (Tun) for the indicated times. Whole-cell extracts were analyzed by western blot with anti-PERK, anti-p-PKR, and anti-PKR antibodies. β-actin was used as a loading control. The analysis was repeated three times, and representative blots from one experiment are shown. **(B) Western blot analysis for p-eIF2α, ATF4, and CHOP in PKR+/+ and PKR−/− MEFs.** Whole-cell extracts from PKR+/+ and PKR−/− MEFs treated the same as in 2A were analyzed at the indicated time points. Blots were probed for p-eIF2α, total eIF2α, ATF4, and CHOP. β-actin was used as a loading control. Representative blots from three independent experiments are shown. **(C) Comparison of the p-eIF2α/total eIF2α ratio in PKR+/+ and PKR−/− MEFs.** The intensities of p-eIF2α and total eIF2α bands were quantified using ImageQuant TL (Cytiva), and the p-eIF2α/eIF2α ratios were calculated. The bar graph shows data from three independent experiments. Black bars: PKR+/+ (PKR wt); white bars: PKR−/− (PKR KO). Error bars represent standard errors. The significant p-values are represented by asterisks.

To assess whether inhibition of PKR with a chemical inhibitor would have a similar effect on ATF4 and CHOP induction, we tested the well-established PKR inhibitor C16 in HeLa cells (Ingrand et al., 2007; Tronel et al., 2014; Szaruga et al., 2023). As seen in Fig. 3A, tunicamycin treatment activated PERK, as evidenced by its upward mobility shift due to phosphorylation at 2-12 h (lanes 3-6), and PKR was activated at 2 h, peaked at 4 h after ER stress (lanes 3-4) and declined at 8 and 12 h post-treatment (lanes 5-6). In the presence of PKR inhibitor C16, PERK activation was observed at 2–12 h after tunicamycin treatment (lanes 9–12), indicating that C16 had no effect on PERK activation at the concentration used, but no PKR activation was seen, as expected (lanes 8–12). These results confirm that C16 inhibits PKR but not PERK activity. We next tested the effect of PKR inhibition on ATF4 and CHOP expression after tunicamycin treatment. As shown in Fig. 3B, in the absence of C16, ATF4 expression was induced 1 h after tunicamycin treatment, steadily increased until 8 h, and declined at 12 h (lanes 2-6). Correspondingly, CHOP expression was seen at 4-12 h after tunicamycin treatment (lanes 4-6). In contrast, in the presence of C16, ATF4 expression was markedly reduced but followed a similar time course, with expression detected at 1 h and increasing until 8 h, then declining at 12 h (lanes 8-12). Correspondingly, CHOP expression was absent at 1-8 h (lanes 8-11), with expression barely detectable at 12 h (lane 12). We further tested the ATF4 and CHOP induction in HeLa DKO cells lacking both PACT and PKR due to targeted CRISPR mutations (Cesaro and Michiels, 2021). As shown in Fig. 3C, both ATF4 and CHOP expression are significantly delayed and reduced in HeLa DKO cells (lanes 8-12) compared with HeLa cells (lanes 2-6). These results further confirm that PKR activation is essential for robust, timely ATF4 and CHOP expression, as seen in murine fibroblasts (Fig. 2), and show that the PKR inhibitor C16 effectively blocks CHOP induction after ER stress.

**Figure 3.**
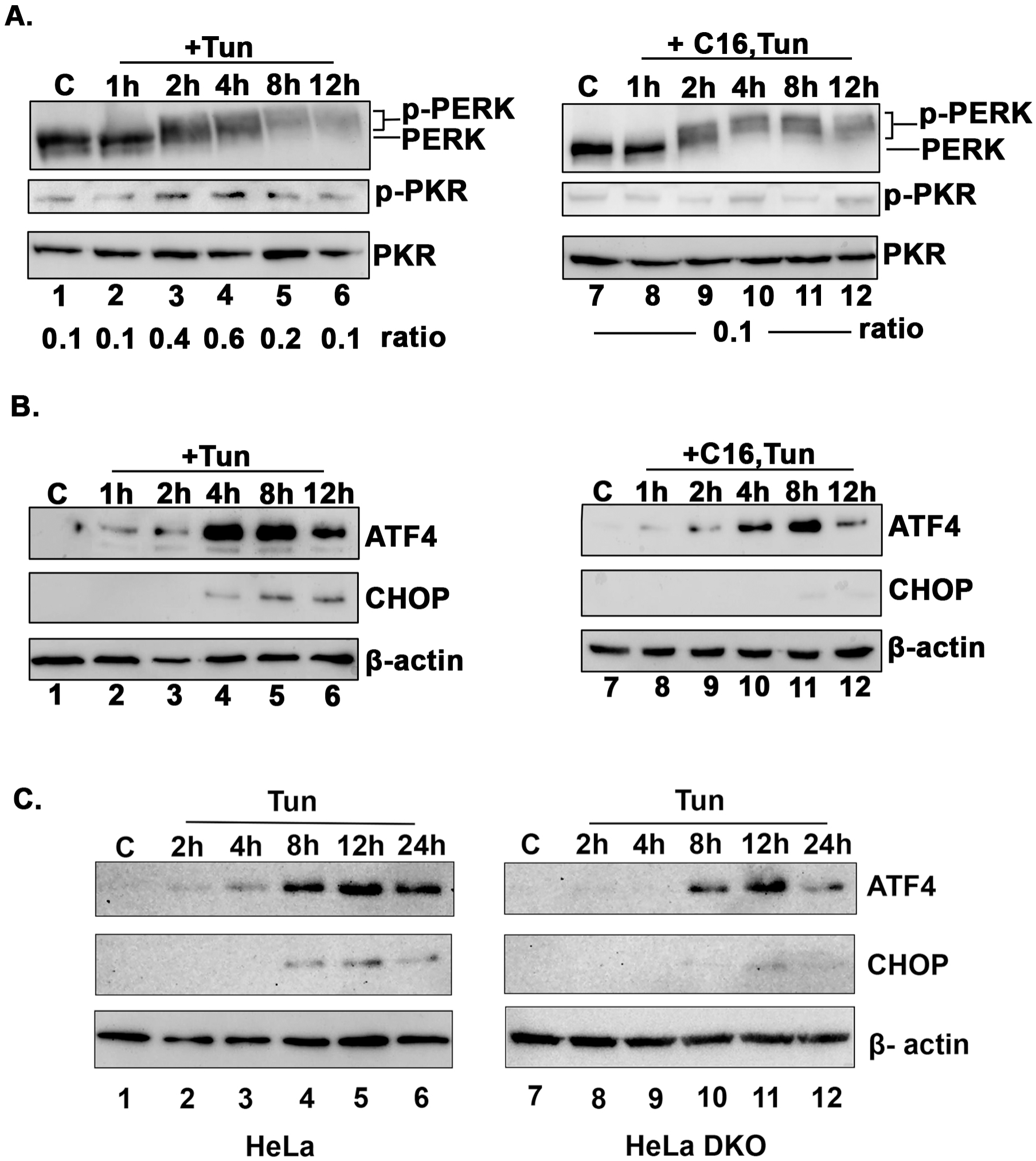
Effect of inhibiting PKR on the response to ER stress in HeLa cells. (A) Effect of the PKR inhibitor C16 on PERK and PKR activation after ER stress. HeLa cells were treated with 0.5 µg/mL tunicamycin (Tun) for the indicated times, either without pre-treatment (+Tun) or after a 2 h pre-treatment with 100 nM C16 (+C16,Tun). Whole-cell extracts were analyzed by western blot with anti-PERK, anti-p-PKR, and anti-PKR antibodies. The ratio of p-PKR to total PKR band intensities is shown below the lanes. The analysis was repeated three times, and representative blots from one experiment are shown. (B) Effect of the PKR inhibitor C16 on ATF4 and CHOP expression after ER stress. HeLa cells were treated the same as in 3A. Whole-cell extracts were analyzed by western blot with anti-ATF4, anti-CHOP, and anti-β-actin antibodies. The analysis was repeated three times, and representative blots from one experiment are shown. (C) Effect of targeted deletion of PKR and PACT on ATF4 and CHOP expression after ER stress. HeLa cells and HeLa DKO cells, which carry targeted inactivation of the PKR and PACT genes, were treated with tunicamycin (Tun) for the indicated times. Whole-cell extracts were analyzed by western blot with anti-ATF4, anti-CHOP, and anti-β-actin antibodies. The analysis was repeated three times, and representative blots from one experiment are shown.

We next tested whether the lack of CHOP expression in the absence of PKR activity results from reduced ATF4-induced CHOP transcription by measuring CHOP mRNA levels by RT-qPCR in HeLa cells treated with tunicamycin in the absence or presence of the PKR inhibitor C16. As seen in Fig. 4A, CHOP mRNA levels were induced about 3-fold at 2 h after tunicamycin treatment and about 13-fold at 4 h after tunicamycin treatment (white bars). In contrast to this, in cells treated with PKR inhibitor C16 prior to tunicamycin treatment, there was no induction of CHOP mRNA at either time point (black bars). These results suggest that the lack of CHOP induction in the absence of PKR activity results from inhibition of ATF4-dependent CHOP transcription after ER stress. We further confirmed this by using a HEK293 ATF4 reporter cell line stably transfected with an ATF4-luciferase reporter construct. In this cell line, ATF4 expression induces luciferase activity, which is directly proportional to the amount of transcriptionally active ATF4. We tested two ER stress inducers: thapsigargin, which induces ER stress by depleting ER calcium stores and inhibiting protein folding (Lytton et al., 1991), and tunicamycin, which inhibits N-linked glycosylation (Heifetz et al., 1979), thereby disrupting protein folding. As seen in Fig. 4B, both thapsigargin and tunicamycin induced luciferase activity about 8-fold and about 15-fold, respectively. A 2 h pre-treatment with PKR inhibitor C16 reduced total luciferase activity by about 50% for thapsigargin and about 80% for tunicamycin. These results further confirm that PKR activity augments ATF4-dependent transcription and is therefore essential for robust CHOP expression. We further investigated PKR’s contribution to the overall ISR using an unbiased RNA-seq approach. RNA-seq analysis was performed on RNA isolated from PKR+/+ and PKR−/− MEFs that were left untreated or treated with tunicamycin for 4 h. As seen in Fig. 4C, an analysis of transcripts that are well-established ATF4 targets (Neill and Masson, 2023) clearly indicates that transcription of these genes is induced robustly in PKR+/+ cells 4 h after tunicamycin treatment but not in PKR−/− cells, where the levels remained close to those in untreated control cells. Transcripts like *Ddit3* (encoding CHOP) and *Ppp1r15a* (encoding GADD34), which were induced robustly at 4 h in PKR+/+ cells, were induced at markedly reduced levels at this time point in PKR−/− cells. Additionally, transcripts encoding several aminoacyl-tRNA synthetases, as well as *Eif2s2* (encoding eIF2β), which are known ATF4 target genes, also showed drastically reduced upregulation at 4 h in PKR−/− cells compared with PKR+/+ cells. These results clearly show that without PKR activity, ATF4 expression and, consequently, transcription of ATF4-regulated genes are drastically reduced. In PKR−/− cells, these transcripts were induced at 12 h after tunicamycin treatment, when their levels began to fall in PKR+/+ cells (data not shown), suggesting a substantial delay in ATF4 induction leading to a corresponding delay in ATF4-induced gene expression. Overall, our results establish that PKR activation during ER stress is required for a timely and efficient ISR and provide heightened and prolonged PKR activation as a plausible reason for increased CHOP expression in DYT-*PRKRA* cells.

**Figure 4.**
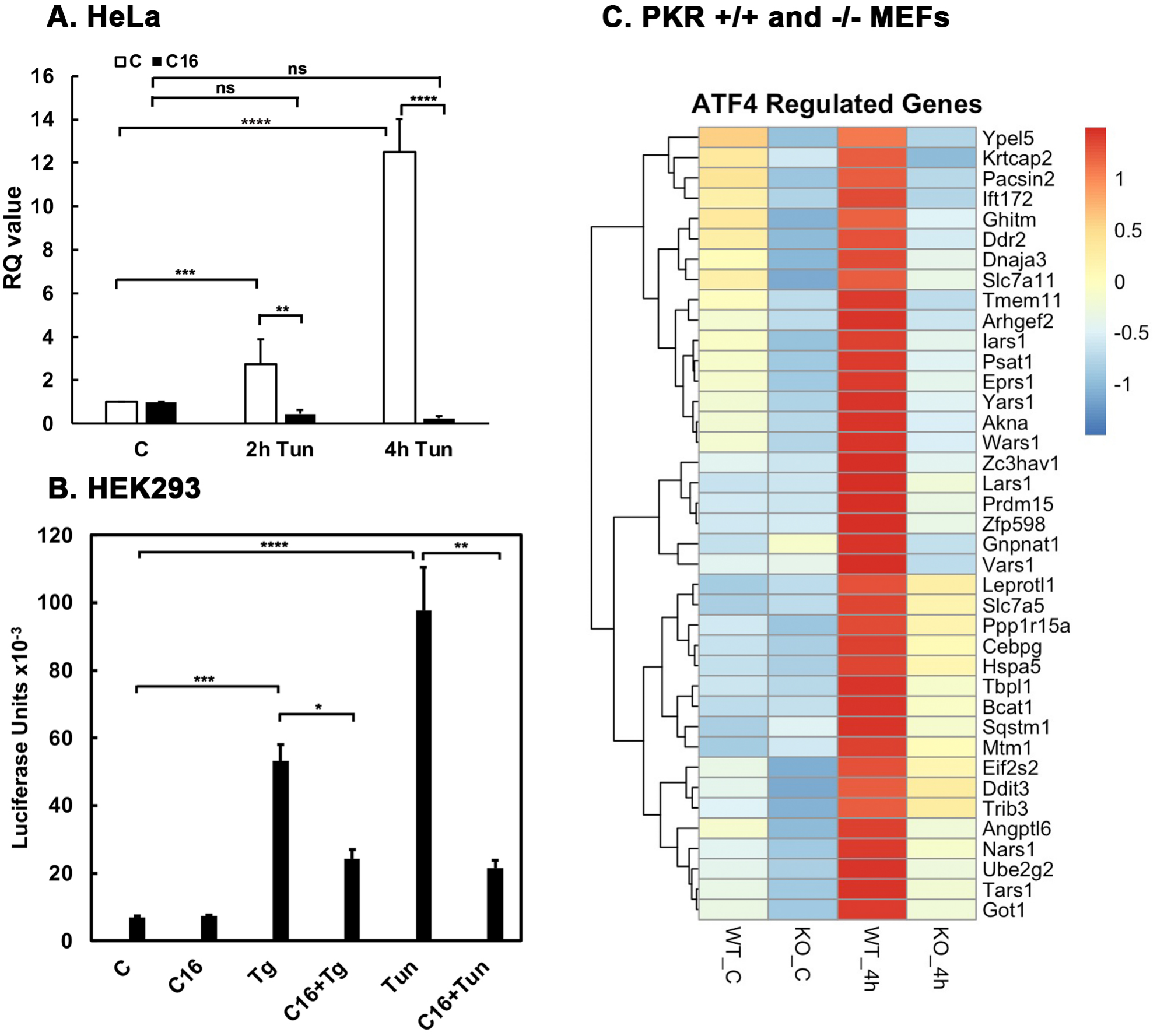
PKR activity is required for ATF4-dependent transcription after ER stress. (A) Effect of PKR inhibition on CHOP mRNA levels analyzed by RT-qPCR. HeLa cells were left untreated (C) or treated with 0.5 µg/mL tunicamycin (Tun) for 2 h or 4 h, with or without a 2 h pre-treatment with 100 nM C16. White bars: without C16; black bars: with C16. CHOP mRNA levels are shown as relative quantification (RQ) values normalized to GAPDH. Error bars represent standard errors. Asterisks indicate that CHOP expression was significantly upregulated at 2 h and 4 h after tunicamycin treatment without C16 compared with control, and was significantly reduced by C16. CHOP induction in the presence of C16 was not significant (ns). **(B) Effect of PKR inhibition on ATF4-dependent transcription in the ATF4-luciferase reporter HEK293 stable cell line.** Cells were plated in triplicate in a 96-well plate and left untreated (C) or treated with 1 µM thapsigargin (Tg) or 0.5 µg/mL tunicamycin (Tun) for 18 h, with or without a 2 h pre-treatment with 100 nM C16. Luciferase activity was measured using a GloMax Explorer plate reader (Promega). The experiment was repeated twice, and the averages were calculated from six replicates. Error bars represent standard errors. The significant p-values are represented by asterisks. **(C) RNA-seq analysis of ATF4 target genes in PKR+/+ and PKR−/− MEFs after tunicamycin treatment.** The heatmap shows the expression of genes known to be upregulated by ATF4 in PKR+/+ (WT) and PKR−/− (KO) MEFs that were left untreated (C) or treated with 0.5 µg/mL tunicamycin for 4 h. Colors show relative expression, from low (blue) to high (red). Data are averaged from three biological replicates.

We next tested whether PKR inhibition can downregulate the increased ATF4 and CHOP induction and apoptosis resulting from excessive PKR activation observed in DYT-*PRKRA* cells, as this would offer a novel treatment strategy for DYT-*PRKRA* and other conditions in which PKR is hyperactive. Our previous results showed that the P222L PACT mutation increases the association of mutant PACT with PKR even without stress and further elevates PKR activation after ER stress (Burnett et al., 2020). The enhanced PKR activation observed in DYT-*PRKRA* lymphoblasts thus results from PACT-mediated PKR activation. Previously, C16 was reported to inhibit PKR when activated by PACT (Farabaugh et al., 2017; Farabaugh et al., 2020; Frederick and Patel, 2023), and we also confirmed this in DYT-*PRKRA* cells in our earlier work. To test whether C16 reduces or blocks tunicamycin-induced apoptosis, we used cleaved PARP1 and caspase activity as apoptosis markers. As seen in Fig. 5A, tunicamycin induced significantly more PARP1 cleavage in DYT-*PRKRA* than in wt lymphoblasts, indicating heightened apoptosis in DYT-*PRKRA* lymphoblasts (compare lanes 2 and 5). Strikingly, PKR inhibition with C16 significantly reduced cleaved PARP1 levels, indicating decreased apoptosis in DYT-*PRKRA* cells (lane 6) as well as in wt lymphoblasts (lane 3). To further confirm that C16 inhibits apoptosis, we used a caspase 3/7 assay. As shown in Fig. 5B, DYT-*PRKRA* patient lymphoblasts (black bars) showed higher caspase activity than wt lymphoblasts (white bars) even without stress, and C16 inhibited this heightened basal caspase activity. At 24 h after tunicamycin treatment, caspase activity increased about 2.2-fold in wt cells and about 1.8-fold in DYT-*PRKRA* cells. C16 inhibited this increase significantly in both wt and DYT-*PRKRA* cells, with about a 40% decrease in wt (white bars) and about a 70% decrease in DYT-*PRKRA* cells (black bars). These results show that PKR inhibition protects both wt and DYT-*PRKRA* cells after ER stress. Finally, to assess the effect of PKR inhibition on ATF4 and CHOP expression during ER stress, we tested the effect of C16 on ATF4 and CHOP after tunicamycin treatment. As seen in Fig. 5C, tunicamycin treatment induced markedly more ATF4 in DYT-*PRKRA* than in wt lymphoblasts at 4 h post-treatment (compare lanes 2 and 5), and C16 blocked ATF4 induction in both wt (lane 3) and DYT-*PRKRA* lymphoblasts (lane 6). CHOP induction was barely detectable at 4 h after ER stress in wt lymphoblasts (lane 2) but was evident in DYT-*PRKRA* lymphoblasts (lane 5), as previously seen in Fig. 1. This induction was completely blocked by C16 in DYT-*PRKRA* lymphoblasts (lane 6), indicating that a plausible reason for the inhibition of apoptosis seen in Fig. 5A and B after C16 treatment is inhibition of CHOP expression. These results indicate that inhibiting PKR can be therapeutically beneficial in dystonia resulting from hyperactive PKR.

**Figure 5.**
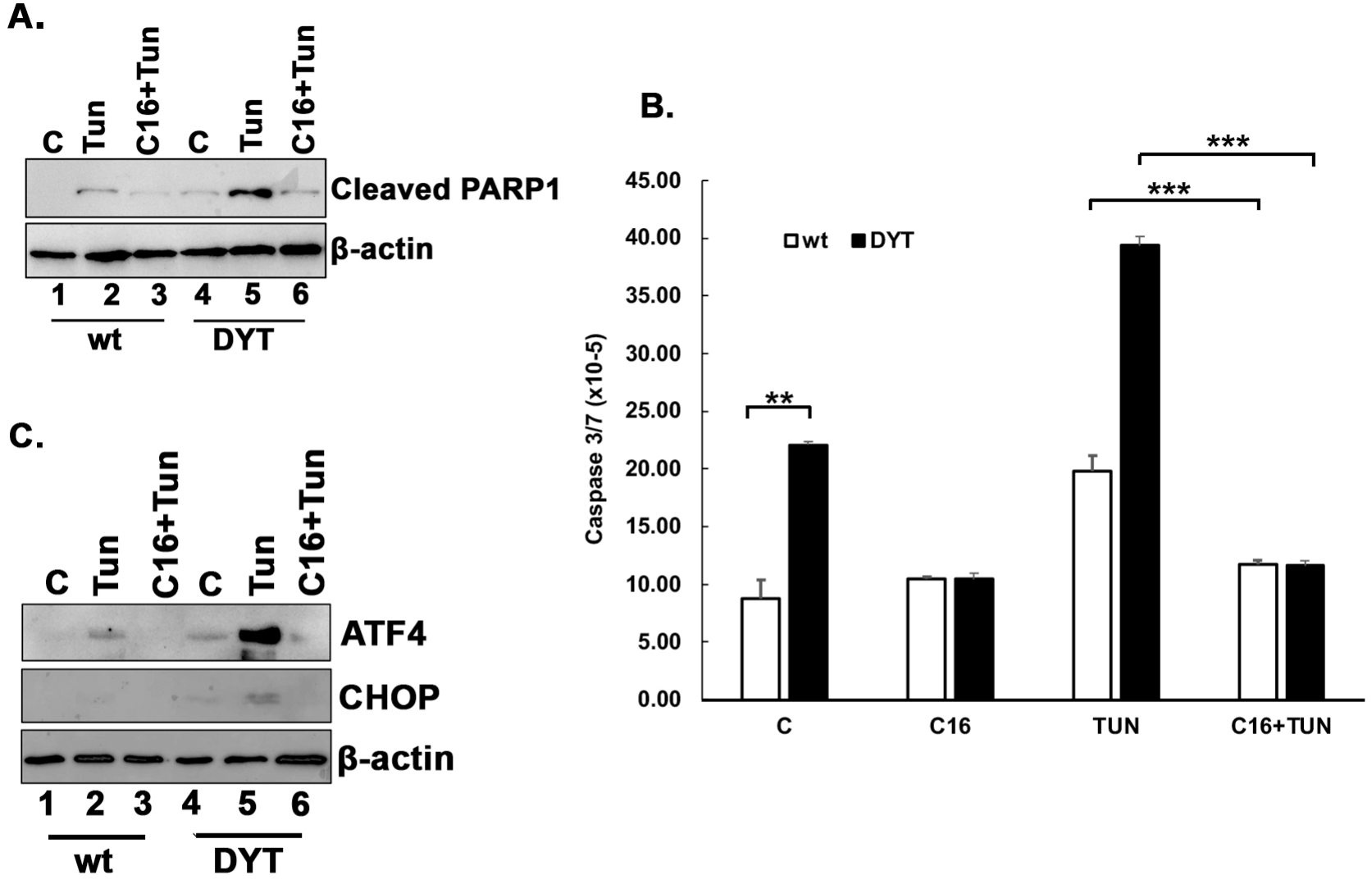
The PKR inhibitor C16 protects DYT-*PRKRA* cells from apoptosis in response to ER stress. (A) Western blot analysis for cleaved PARP1. Normal (wt) and DYT-*PRKRA* (DYT) patient-derived lymphoblasts were left untreated (C) or treated with 0.5 µg/mL tunicamycin (Tun), with or without a 2 h pre-treatment with 100 nM C16 (C16+Tun). Whole-cell extracts prepared 24 h after tunicamycin treatment were analyzed by western blot with anti-cleaved PARP1 and anti-β-actin antibodies. β-actin was used as a loading control. **(B) Caspase 3/7 activity.** Normal (wt) and DYT-*PRKRA* (DYT) patient-derived lymphoblasts were left untreated (C), treated with 100 nM C16 alone (C16), treated with 0.5 µg/mL tunicamycin (Tun), or pre-treated with 100 nM C16 for 2 h before tunicamycin treatment (C16+Tun). Caspase 3/7 activity was measured at 24 h using the Caspase-Glo 3/7 assay. White bars: wt cells; black bars: DYT-*PRKRA* cells. The data are the average of three independent experiments, and error bars represent standard errors. The significant p-values are represented by asterisks. **(C) Western blot analysis for ATF4 and CHOP.** Normal (wt) and DYT-*PRKRA* (DYT) patient-derived lymphoblasts were treated the same as in 5A, and Whole-cell extracts were prepared 4 h after tunicamycin treatment. Extracts were analyzed by western blot with anti-ATF4, anti-CHOP, and anti-β-actin antibodies. β-actin was used as a loading control. Blots in (A) and (C) are representative of three independent experiments.

## Discussion

DYT-*PRKRA* is a debilitating, early-onset movement disorder for which there are currently no effective treatment options. Although most DYT-*PRKRA* cases have early childhood onset, later onset in adulthood has also been observed, suggesting environmental or other modifying genetic factors (Camargos et al., 2008; Zech et al., 2014). A few patients show abnormalities and degeneration in the striatal region, but this information was not available for most patients (Kolbel et al., 2017; Masnada et al., 2021; Bhowmick et al., 2022). Our results suggest that increased apoptosis in DYT-*PRKRA* cells, driven by heightened PKR activation, could contribute to the observed striatal degeneration, although this remains to be tested in neuronal models. Notably, in several cases, dystonia symptoms were first noted or often worsened after a childhood febrile illness (Lemmon et al., 2013; Kolbel et al., 2017; Pinto et al., 2020; Masnada et al., 2021; Bhowmick et al., 2022). Viral infections cause ER stress because they require large amounts of viral structural proteins to be synthesized relatively quickly (Zhang and Wang, 2012; Chen et al., 2026). Additionally, PKR is also activated during viral replication independently of ER stress, by viral nucleic acids and replication intermediates (Dauber and Wolff, 2009). In febrile illness episodes, although not necessarily viral infections, excessive PKR activation, as observed in our DYT-*PRKRA* cell culture studies, could worsen movement abnormalities. Currently, there are no therapeutic drugs available for DYT-*PRKRA*, and the frequently used deep brain stimulation (DBS) therapeutic intervention for several types of dystonia has not shown much benefit in DYT-*PRKRA* patients (Tisch and Kumar, 2020) although benefit of DBS has been reported recently in three patients (Gomes et al., 2026). Other established treatments, including botulinum toxin injections, baclofen, and benzodiazepines, have also been shown not to be beneficial (Thomsen et al., 2024). Thus, understanding the molecular mechanisms underlying DYT-*PRKRA* can be considered a top priority for identifying novel and effective drug targets.

At least 17 distinct *PRKRA* mutations have been reported in DYT-*PRKRA* patients, including recessive and dominant missense variants, compound heterozygous alleles, a noncoding variant, and a frameshift mutation (Simon and Patel, 2025). Among the reported disease alleles, the homozygous *PRKRA* c.665C>T (p.Pro222Leu; P222L) variant is the most frequently reported mutation in the published DYT-*PRKRA* literature and is represented in the original Brazilian families as well as in affected individuals from Italy, Portugal, Spain, and Turkey (Simon and Patel, 2025; Gomes et al., 2026). P222L is therefore the best-characterized DYT-*PRKRA* mutation clinically and mechanistically. Our previous research showed that many DYT-*PRKRA* variants lead to enhanced PKR activation and eIF2α phosphorylation in response to ER stress. The recessively inherited P222L variant causes delayed but heightened and persistent PKR activation and eIF2α phosphorylation, increasing cell susceptibility to ER stress (Vaughn et al., 2015). A dominantly inherited frameshift variant causes insoluble protein aggregates, PKR activation, and significant apoptosis even without stress (Burnett et al., 2019). Additionally, three other recessive and two dominant variants also increase PKR interaction and activation, causing increased eIF2α phosphorylation and sensitivity to ER stress in DYT-*PRKRA* patient cells (Burnett et al., 2020). Understanding the molecular mechanisms driving enhanced apoptosis in cell culture models of DYT-*PRKRA* is important for designing effective, mechanistically grounded treatment strategies.

Previously, based on our discovery that the plant flavonoid luteolin disrupts the interaction between PACT and PKR in human cells in culture, we tested it as a therapeutic option for DYT-*PRKRA* (Frederick and Patel, 2023). Luteolin effectively disrupted the enhanced interactions between PKR and the P222L mutant, protecting DYT-*PRKRA* patient cells from increased apoptosis in response to ER stress. Although promising, luteolin is not specific to PACT-PKR interactions and has pleiotropic effects across several cellular pathways (Caporali et al., 2022; Wang et al., 2025). A drug that specifically targets PKR to inhibit its excessive catalytic activity in DYT-*PRKRA* cells is better suited for clinical use. Notably, our results show that PKR inhibition with C16 reduces the expression of both ATF4 and CHOP in DYT-*PRKRA* cells, thereby establishing PKR inhibition as a potential therapeutic option.

Although our previous studies described enhanced PKR activation after ER stress in DYT-*PRKRA* patient cells carrying the homozygous P222L mutation, this study is the first to delineate downstream molecular events, such as ATF4 and CHOP induction, following PKR activation and eIF2α phosphorylation. Importantly, our results indicate that PERK activation in response to ER stress is unaffected by the P222L mutant and establish that PERK does not contribute to the delayed and persistent eIF2 phosphorylation observed in DYT-*PRKRA* cells. The persistent increase in eIF2α phosphorylation in DYT-*PRKRA* thus results solely from heightened and persistent PKR activity. The corresponding increased and persistent expression of ATF4 and CHOP observed in DYT-*PRKRA* cells also results solely from dysregulated PKR activation. Although PKR is a well-characterized eIF2α kinase, it is mainly studied in the context of viral infections (Samuel et al., 1997; Chukwurah et al., 2021) and is activated by binding to double-stranded (ds) RNA often produced during viral replication (Cole, 2007). PERK and PKR are thus canonically thought to respond to distinct upstream stress signals, with PERK mainly responding to ER stress and PKR mainly responding to stress caused by viral infections. However, many studies have indicated that PERK and PKR respond to overlapping sets of stress signals, sometimes with crosstalk during viral infections as well as several other conditions that elicit ER stress (Yan et al., 2002; Baltzis et al., 2004). Our results show that PKR activation plays a pivotal role in timely, robust ATF4 induction and the subsequent downstream transcriptional events regulated by ATF4. Without PKR activation, ATF4 induction is significantly delayed and reduced. Consequently, ATF4-regulated gene expression is also compromised, as shown by our RNA-seq analysis. Our results thus establish for the first time that in addition to PERK activation after ER stress, PKR activation is essential for an effective and timely expression of ATF4-regulated genes.

There can be various reasons for activation of two eIF2α kinases (PERK and PKR) when cells experience ER stress. PERK is embedded in the ER membrane, and its cytoplasmic catalytic domain phosphorylates eIF2α (Volchuk and Ron, 2010). PERK activation by ER stress involves trans-autophosphorylation, which leads to phosphorylation of multiple serine and threonine residues in its unusually long kinase insert loop (Marciniak et al., 2006). Using elegant biochemical experiments, Marciniak et al. showed that PERK activation and phosphorylation selectively increase its affinity for the non-phosphorylated eIF2 complex because ER stress activation induces a major conformational change in the PERK kinase domain. Additionally, they also demonstrated that although dispensable for catalytic activity, PERK’s kinase insert loop is required for substrate binding and for eIF2α phosphorylation *in vivo*. This offered a mechanism for eIF2 recruitment by activated PERK and for unidirectional substrate flow, where phosphorylated eIF2 dissociated and freely diffused in the cytoplasm to interact with eIF2B and exert its inhibitory actions to halt protein synthesis (Marciniak et al., 2006). Nonetheless, because PERK is membrane-bound, it may be insufficient to phosphorylate a significant fraction of eIF2 to produce an efficient, timely block in protein synthesis. Our results support this idea, as in the absence of PKR the eIF2a phosphorylation levels are lower and the resulting block in protein synthesis is also significantly delayed (data not shown). Consequently, ATF4 induction is markedly reduced and delayed, with ATF4-dependent CHOP induction being barely detectable. At later time points, cells eventually catch up with ATF4 and CHOP induction in the absence of PKR (data not shown), indicating that PKR significantly increases both the speed and level of eIF2a phosphorylation in response to ER stress.

The essential function of PKR in supplementing and augmenting the PERK-initiated ER stress response is further supported by our data demonstrating markedly reduced ATF4-luciferase reporter activity in HEK293 cells when PKR is inhibited using its small-molecule inhibitor C16 (Fig. 4). These results indicate that ATF4 is not fully active as a transcription factor in the absence of PKR activity. ATF4 is a central regulator of cellular fate after ER stress, and the ATF4-dependent downstream pathways are involved in regulating pro-survival or pro-apoptotic outcomes (Pitale et al., 2017; Neill and Masson, 2023). Combinations of ATF4’s heterodimerization partners and post-translational modifications are known to influence this regulation (Neill and Masson, 2023). In the absence of PKR activity, ATF4-dependent transcription is markedly delayed and reduced; PKR may directly or indirectly control ATF4 post-translational modifications, which could influence its transcriptional activation potential by determining its dimerization partners and affecting expression of ATF4 target genes. Additionally, PKR activity may be essential for the IRE1 and/or ATF6 pathways activated after ER stress, which are known to co-regulate ATF4’s transcriptional targets (Schroder and Kaufman, 2005). Our analysis of known ATF4-regulated genes from a large RNA-seq dataset of tunicamycin-treated PKR+/+ and PKR−/− MEFs showed significantly reduced expression of ATF4 target genes in PKR−/− cells at 4 h after treatment. These results confirm the western blot and RT-qPCR analyses of CHOP expression (Fig. 2, 3 and 4A) and suggest that the delay in ATF4-dependent gene expression in the absence of PKR applies to a much larger set of genes. Future detailed analysis of this large dataset is likely to reveal additional genes that require PKR activity for transcriptional induction after ER stress. Fig. 6 summarizes our main findings that are relevant to ISR regulation as well as DYT-*PRKRA*.

**Figure 6.**
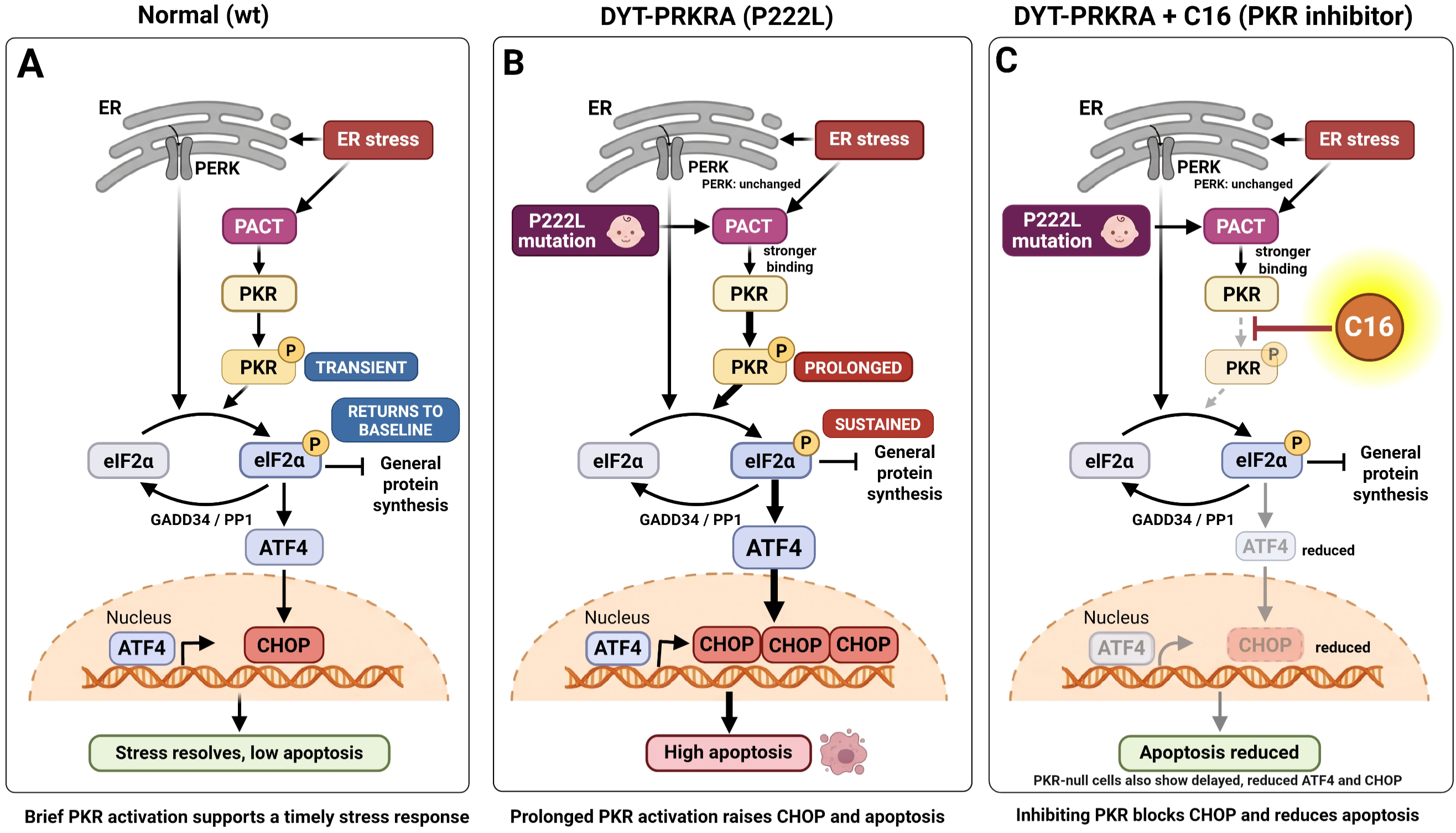
Model of the PKR-dependent response to ER stress in normal and DYT-*PRKRA* cells, and the effect of PKR inhibition. (A) Normal (wt) cells. ER stress activates PERK and, separately, activates PKR through PACT (Fig. 1A). Both kinases phosphorylate eIF2α (P), which inhibits general protein synthesis and induces ATF4. PKR activation is transient and eIF2α phosphorylation returns to baseline (Fig. 1A, B), in part through GADD34/PP1-mediated dephosphorylation. ATF4 induces a low level of CHOP (Fig. 1C, D), the stress resolves; apoptosis is low (Fig. 5A, B), and the cell returns to homeostasis. **(B) DYT-*PRKRA* (P222L) cells.** The P222L mutation strengthens the PACT–PKR interaction. PERK activation is unchanged in DYT-*PRKRA* cells (Fig. 1A), but PKR activation is prolonged, and eIF2α phosphorylation is sustained (Fig. 1A, B), leading to higher ATF4 and CHOP levels (Fig. 1C, D) and high apoptosis (Fig. 5A, B). Thick arrows indicate the enhanced steps. **(C) DYT-*PRKRA* cells treated with the PKR inhibitor C16.** C16 blocks PKR activation without affecting PERK phosphorylation (Fig. 3A). ATF4 and CHOP induction are reduced (Fig. 5C), and apoptosis is reduced (Fig. 5A, B). PKR-null cells show a similar delay and reduction in ATF4 and CHOP induction (Fig. 2B). Dashed arrows and faded shapes indicate steps that are blocked or reduced. The highlight marks the site of C16 action. Created in BioRender. Patel, R. (2027) https://BioRender.com/g3yjadb

While it is exciting that PKR inhibition with its specific chemical inhibitor C16 protects DYT-*PRKRA* cells, several critical points must be considered for therapeutic use when regulating ISR via PKR inhibition. Other than C16, 2-aminopurine (2-AP) is another small molecule inhibitor of PKR that has been characterized (Hu and Conway, 1993). Both inhibitors act on the ATP-binding site of PKR and have limitations. C16 has a low IC50 but is not specific for PKR (Chen et al., 2008), and 2-AP has a high IC50 and low specificity for PKR (Huang et al., 2003). Importantly, our conclusions do not rest on C16 alone, because PKR^−/−^ MEFs and HeLa DKO cells showed a similar delay and reduction in ATF4 and CHOP induction. C16 has limited bioavailability, making it difficult to maintain a stable, effective concentration in human tissues. As PKR plays a central role in innate antiviral response, pharmacological inhibition of PKR could impair the primary antiviral state, leaving individuals highly susceptible to severe or lethal viral infections. Additionally, PKR is important for adaptive immune responses, regulating T cell activation and production of specific inflammatory cytokines, and inflammasome activation (Chukwurah et al., 2021). Consequently, utilizing therapies that inhibit PKR could severely undermine these vital host defense mechanisms. Future studies could explore a drug that targets only the enzymatically active conformation of PKR, which seems plausible since active PKR differs conformationally from catalytically inactive PKR.

Dysregulated ISR has emerged as a convergent mechanism in various monogenic dystonia types (Gonzalez-Latapi et al., 2021; Calakos and Caffall, 2024; Calakos and Zech, 2025). In contrast to the excessive PKR activation and eIF2α phosphorylation in DYT-*PRKRA* patient cells, basal PKR activity and eIF2α phosphorylation levels are reduced in *PRKRA^lear-5J^* mice (Burnett et al., 2024), suggesting that reduced PKR activity and, consequently, reduced eIF2α phosphorylation may also contribute to dystonia pathophysiology. Several mutations in the *EIF2AK2* gene, which encodes PKR, have been reported to cause PKR hyperactivation, upregulated eIF2α phosphorylation, and childhood-onset DYT-*EIF2AK2* (DYT33) (Kuipers et al., 2021). Some *EIF2AK2* mutations that inactivated or reduced PKR activity were also reported to cause a broad spectrum of neurodevelopmental abnormalities, including developmental delays, variable cognitive impairments, white matter alterations, and dystonia (Mao et al., 2020). Thus, in PKR-mediated regulation of eIF2α phosphorylation levels, both lower- and higher-than-optimal levels are detrimental to coordinated muscle movements. One missense *EIF2AK1* (HRI) mutation with reduced eIF2α phosphorylation and ISR dysregulation was also reported to cause movement abnormalities (Mao et al., 2020). DYT-*TOR1A*, is a childhood-onset autosomal-dominant disease caused by a single amino acid deletion in the ER-resident torsinA protein. DYT-*TOR1A* patient cells exhibit an activated ER stress response, and eIF2α signaling is dysregulated (Rittiner et al., 2016). Remarkably, in DYT-*TOR1A*, restoring eIF2α phosphorylation to normal levels alleviated dystonia symptoms (Caffall et al., 2021). DYT-*THAP1* is caused by mutations in *THAP1* (Zakirova et al., 2018), and a transcriptomic analysis in the neonatal mouse striatum and cerebellum indicated dysregulation of the eIF2α signaling pathway; salubrinal, which promotes eIF2α phosphorylation, partially corrected neuronal plasticity defects (Boyce et al., 2005). DYT-*SGCE* is caused by mutations in the *SGCE* gene, which encodes ε-sarcoglycan (ε-SG), and PKR is upregulated in a DYT-*SGCE* mouse model (Xiao et al., 2017). Strikingly, several mutations in *ATF4* itself have been reported in patients with sporadic cervical dystonia (Rittiner et al., 2016). Additionally, several dystonia genes related to calcium physiology also have altered ISR and eIF2α signaling (Jinnah and Sun, 2019).

Similar to PKR and eIF2α phosphorylation, there is also evidence for both high and low activity levels of ATF4 being detrimental to neurons. Under physiological conditions, baseline ATF4 levels support synaptic plasticity and neuronal maturation, promoting homeostasis, while insufficient or chronically low ATF4 activity impairs adaptive responses to metabolic and cellular stress, compromising neuronal differentiation and resilience (Sun et al., 2013). Conversely, sustained or pathologically high ATF4 expression triggers pro-apoptotic signaling cascades and promotes neuronal death (Chen et al., 2017). This dual, context-dependent nature underscores an inverted U-shaped relationship between ATF4 dosage and neuronal viability. Excessively low ATF4 activity leaves neurons vulnerable to environmental insults by disrupting protein homeostasis. Conversely, ATF4 hyperactivation transcriptionally upregulates death-associated target genes, accelerating neurodegeneration (Demmings et al., 2021; Demmings et al., 2026). Beyond classical neurodegenerative disorders, dysregulated ATF4 signaling is also directly implicated in motor circuit dysfunction. In particular, both sub- and supra-optimal ATF4 signaling disrupt striatal neuromodulation and synaptic connectivity, provoking hyperkinetic posturing and motor deficits characteristic of DYT-*TOR1A* (Calakos and Caffall, 2024). Our DYT-*PRKRA* results highlight how both insufficient stress adaptability and persistent ATF4/CHOP-mediated apoptotic induction contribute to striatal circuit vulnerability and dystonia pathogenesis. Consequently, maintaining a precise, optimal homeostatic window of ATF4 expression is imperative to safeguard neuronal integrity and motor circuit function against toxicities resulting from both insufficiency and overactivation. Any therapeutic development would thus need to be fine-tuned to the specific molecular details of the condition being treated.

Beyond its role as a stress response pathway, the ISR contributes to physiological processes in the nervous system, including synaptic plasticity, memory formation, axon development, and circadian rhythms (Lockshin and Calakos, 2024). Regulated eIF2α phosphorylation controls the translation of proteins required for various neuronal functions, and genetic or pharmacological manipulations that alter eIF2α phosphorylation levels or duration impair neuronal function (Lockshin and Calakos, 2024). In specific neuronal populations, such as striatal cholinergic interneurons (CINs), the ISR is constitutively active and inhibiting the ISR in CINs of normal mice reproduces the inverted physiological response to dopamine type 2 receptor (D2R) agonists characteristic of dystonia mouse models (Helseth et al., 2021), By regulating the mRNAs or ORFs selectively translated through eIF2α phosphorylation, ISR strongly influences the neuronal proteome and, consequently, basic functions such as muscle movement and coordination that are not linked to stress signaling. This neuronal sensitivity means that dysregulation of ISR signaling, either through excessive or impaired activation, or through alterations in its optimal physiological duration, can have particularly profound consequences in the nervous system (Calakos and Caffall, 2024). Not surprisingly, ISR dysregulation is associated with many neurodegenerative and neurodevelopmental diseases, including but not limited to age-related cognitive disorders, intellectual disability, Alzheimer’s disease, Parkinson’s disease, frontotemporal dementia, Down syndrome, and prion diseases (Costa-Mattioli and Walter, 2020). Specifically, excessive PKR activation is noted in several of these conditions, and our results strongly indicate that PKR is an important drug target that has remained underexplored and is promising for future drug development for several pathologies in addition to DYT-*PRKRA*.

## Materials and Methods

### Reagents, Cell Lines, and Antibodies

The wild-type (wt; normal) and DYT-*PRKRA* patient B-lymphoblast cell lines, established by Epstein-Barr virus transformation as previously described (Anderson and Gusella, 1984) were cultured in RPMI 1640 medium supplemented with 10% fetal bovine serum (FBS) and penicillin/streptomycin. The lymphoblast lines were a generous gift from Dr. Bragg and Sharma (Dystonia Partners Research Bank, Massachusetts General Hospital, USA). HeLa cells, HeLa DKO cells and PKR+/+ and PKR-/- mouse embryonic fibroblasts (MEFs) were cultured in Dulbecco’s Modified Eagle’s Medium (DMEM) supplemented with 10% FBS and penicillin/streptomycin. Tunicamycin (sc-203119C) was purchased from Santa Cruz Biotechnology. Thapsigargin (586005) and the PKR inhibitor C16 (527450) were purchased from Sigma-Aldrich. The following primary antibodies were used: **PERK:** anti-PERK (Abcam, ab229912), **PKR:** anti-human PKR (R&D Systems, MAB1980), **PKR:** anti-mouse PKR (Abcam, ab184257), **Phospho-PKR:** anti-phospho-PKR (Thr446) (Abcam, ab32036), **eIF2α**: anti-eIF2α (Cell Signaling Technology, 9722S), **Phospho-eIF2α**: anti-phospho-eIF2α (Ser51) (Cell Signaling Technology, 3398S), **ATF4:** anti-ATF4 (Cell Signaling Technology, 11815S), **CHOP:** anti-CHOP (Cell Signaling Technology, 2895S), **Cleaved PARP1:** anti-cleaved PARP1 (Cell Signaling Technology, 5625S), and peroxidase-conjugated anti-β-actin (Sigma-Aldrich, A3854). HRP-conjugated secondary antibodies were goat anti-mouse (Invitrogen, 62-6520) and goat anti-rabbit (Cell Signaling Technology, 7074P2).

### Western Blot Analysis

Human lymphoblast lines, HeLa cells, HeLa DKO cells, and PKR+/+ and PKR-/- MEFs were treated with 0.5 µg/mL tunicamycin. All cells were harvested at the indicated time. PKR activity was inhibited using 100 nM C16 for 2 h before ER stress induction. Cells were washed twice with ice-cold phosphate-buffered saline (PBS) and lysed in RIPA buffer. Protein concentrations were determined using a BCA assay. Equal amounts of protein lysates were resolved using sodium dodecyl sulfate-polyacrylamide gel electrophoresis (SDS-PAGE) and transferred onto polyvinylidene difluoride (PVDF) membranes. The membranes were blocked with either 5% milk or 5% bovine serum albumin (BSA) for 1 h and incubated with primary antibodies overnight at 4°C, followed by incubation with HRP-conjugated secondary antibodies for 1 h at room temperature. Chemiluminescence was detected using the ImageQuant LAS 4000 system (Amersham). Band intensities were quantified using ImageQuant TL software (Cytiva).

### Caspase 3/7 Assay

Patient and wt lymphoblasts were plated at 1 × 10^5^ cells/well in 0.1 mL in a 96-well cell culture plate and were left untreated, treated with 100 nM C16 alone, treated with 0.5 µg/mL tunicamycin, or pre-treated with 100 nM C16 for 2 h before tunicamycin treatment. At 24 h after tunicamycin treatment, the cells were mixed with an equal volume of Caspase-Glo 3/7 reagent (Promega) and incubated for 30 min. Luminescence was measured using a GloMax Explorer plate reader (Promega). Wells containing cell culture medium alone served as the negative control for background luminescence and were used to normalize all readings.

### ATF4-luciferase Reporter Assay

The ATF4 Luciferase Reporter HEK293 Stable Cell Line (Signosis, SL-0080) is derived from human embryonic kidney cells and stably expresses a firefly luciferase reporter gene under the control of the ATF4 DNA-binding response element. The cells were grown in DMEM supplemented with 10% fetal bovine serum (FBS), penicillin/streptomycin, and 50 µg/mL hygromycin B to maintain the selection pressure. Cells were plated at 20,000 cells per well in 0.1 mL in a 96-well plate and were left untreated or treated with 1 µM thapsigargin or 0.5 µg/mL tunicamycin, with or without a 2 h pre-treatment with 100 nM C16. At 18 h after treatment, an equal volume of ONE-Glo reagent (Promega) was added to the wells, and luciferase activity was measured after 5 min using a GloMax Explorer plate reader (Promega).

### RNA Extraction and Quantitative RT-PCR (RT-qPCR)

HeLa cells were treated with 0.5 µg/mL tunicamycin for the indicated time points. In the samples where PKR was inhibited, cells were pre-treated with 100 nM C16 (PKR inhibitor) for 2 h before tunicamycin treatment. Total RNA was extracted using TRIzol reagent (Thermo Fisher Scientific, 15596026) following the manufacturer’s protocol. RNA concentration and purity were assessed by spectrophotometry (NanoDrop). Complementary DNA (cDNA) synthesis was performed using SuperScript IV Reverse Transcriptase (Thermo Fisher Scientific, 18090010) according to the manufacturer’s instructions. Quantitative PCR was performed using GoTaq® Hot Start DNA Polymerase (Promega, M7401) in combination with EvaGreen® dye (Biotium, 31019) for fluorescence detection. The expression levels of the target genes were normalized to GAPDH, and relative gene expression was determined using the ΔΔCt method. Primer sequences were as follows: **CHOP:** Forward: 5’-GTT AAA GAT GAG CGG GTG GC-3’, Reverse: 5’-GCT TTC AGG TGT GGT GAT GTA T-3’, **GAPDH** (housekeeping control): Forward: 5’-CAT GGG TGG AAT CAT ATT GGA AC-3’, Reverse: 5’-GGT CGG AGT CAA CGG ATT TGG-3’.

### RNA-seq Analysis

To induce ER stress, PKR+/+ and PKR−/− MEFs at 50–60% confluency were left untreated or treated with 0.5 µg/mL tunicamycin for 4 h or 12 h. Three independent biological replicates were collected for each condition and genotype. Total RNA was isolated using the Direct-zol RNA MiniPrep Plus Kit (Zymo Research, R2070) according to the manufacturer’s instructions. RNA concentration and purity were determined by spectrophotometry (NanoDrop), and RNA integrity was assessed using an Agilent 2100 Bioanalyzer. Only samples with an RNA integrity number (RIN) ≥ 7.0 were used for sequencing. RNA samples were submitted to Novogene (Sacramento, CA, USA) for library preparation and next-generation sequencing. Library construction, quality control, and sequencing were performed by the service provider using standard Illumina-compatible protocols. Briefly, mRNA was enriched by poly(A) selection, fragmented, and converted to cDNA prior to adapter ligation and amplification. The Sequencing was performed on an Illumina NovaSeq X Plus to generate 150 bp paired-end reads. Raw sequencing data (FASTQ files) and BAM files of reads aligned to the mouse reference genome (mm39) were returned to the authors for downstream analysis.

### RNA-seq Data Analysis

Differentially expressed genes (DEGs) were identified using the *DESeq2* package in R, with n = 3 biological replicates per condition. A false discovery rate (FDR) threshold of < 0.05 was applied to determine statistical significance. To visualize the expression profiles of known ATF4 downstream targets, hierarchical clustering heatmaps were generated using the *pheatmap* package in R. (Kolde R. https://cran.r-project.org/web/packages/pheatmap/index.html).

### Statistical Analysis

The western blot band intensity data were analyzed using Microsoft Excel, and error bars are presented as means ± standard error of the mean (SEM). To determine statistically significant differences between groups, we performed two-tailed Student’s t-tests. Statistical significance was defined a priori at a threshold of *p* < 0.05. In figures, significance is denoted as *p < 0.05, **p < 0.01, ***p < 0.001, ****p < 0.0001.

## Acknowledgements

We thank Tieqiang Hu for technical help with RT-qPCR analysis. We also thank Dr. Cristopher Bragg and Dr. Nutan Sharma for normal and patient DYT-*PRKRA* lymphoblast lines and Dr. Bryan G. Williams for PKR +/+ and -/- MEFs. We thank Sanjana Parise for expert technical help with western blots.

## Author contributions

E. G. M. and S. A. designed and performed experiments, interpreted results, analyzed data, and wrote the manuscript; T. R. performed the HeLa and C16 experiments, J. C. A. performed the HeLa DKO experiment, and both analyzed their respective data and edited the manuscript. R. C. P. conceptualized and supervised the work, acquired funding, and edited the manuscript.

## Data and resource availability

All cell lines, plasmids, and other resources used in this study will be made available by the authors upon reasonable request. The RNA-seq datasets supporting the conclusions of this article will be deposited in NCBI GEO prior to publication, and the accession number/link will be provided.

## References

Adjibade, P. and Mazroui, R. (2026) Regulation of Translation of ATF4 mRNA: A Focus on Translation Initiation Factors and RNA-Binding Proteins, Cells 15(2).

Albanese, A., Bhatia, K., Bressman, S. B., Delong, M. R., Fahn, S., Fung, V. S., Hallett, M., Jankovic, J., Jinnah, H. A., Klein, C. et al. (2013) Phenomenology and classification of dystonia: a consensus update, Movement Disorders 28(7): 863–873.

Anderson, M. A. and Gusella, J. F. (1984) Use of cyclosporin A in establishing Epstein-Barr virus-transformed human lymphoblastoid cell lines, In Vitro 20(11): 856–858.

Atasu, B., Simón-Sánchez, J., Hanagasi, H., Bilgic, B., Hauser, A. K., Guven, G., Heutink, P., Gasser, T. and Lohmann, E. (2024) Dissecting genetic architecture of rare dystonia: genetic, molecular and clinical insights, Journal of Medical Genetics 61(5): 443–451.

Baltzis, D., Qu, L. K., Papadopoulou, S., Blais, J. D., Bell, J. C., Sonenberg, N. and Koromilas, A. E. (2004) Resistance to vesicular stomatitis virus infection requires a functional cross talk between the eukaryotic translation initiation factor 2alpha kinases PERK and PKR, Journal of Virology 78(23): 12747–12761.

Bennett, R. L., Blalock, W. L., Abtahi, D. M., Pan, Y., Moyer, S. A. and May, W. S. (2006) RAX, the PKR activator, sensitizes cells to inflammatory cytokines, serum withdrawal, chemotherapy, and viral infection, Blood 108(3): 821–829.

Bennett, R. L., Blalock, W. L. and May, W. S. (2004) Serine 18 phosphorylation of RAX, the PKR activator, is required for PKR activation and consequent translation inhibition, Journal of Biological Chemistry 279(41): 42687–42693.

Bhowmick, S. S., Raha, S. and Bohora, A. (2022) Early-Onset Dystonia, Exacerbation With Fever, and Striatal Signal Changes: Emerging Phenotype of DYT-PRKRA, Neurology 99(5): 206–207.

Boyce, M., Bryant, K. F., Jousse, C., Long, K., Harding, H. P., Scheuner, D., Kaufman, R. J., Ma, D., Coen, D. M., Ron, D. et al. (2005) A selective inhibitor of eIF2alpha dephosphorylation protects cells from ER stress, Science 307(5711): 935–939.

Bragg, D. C., Armata, I. A., Nery, F. C., Breakefield, X. O. and Sharma, N. (2011) Molecular pathways in dystonia, Neurobiology of Disease 42(2): 136–147.

Burnett, S. B., Culver, A. M., Simon, T. A., Rowson, T., Frederick, K., Palmer, K., Murray, S. A., Davis, S. W. and Patel, R. C. (2024) Mutation in Prkra results in cerebellar abnormality and reduced eIF2α phosphorylation in a model of DYT-PRKRA, Disease Models & Mechanisms 17(11).

Burnett, S. B., Vaughn, L. S., Sharma, N., Kulkarni, R. and Patel, R. C. (2020) Dystonia 16 (DYT16) mutations in PACT cause dysregulated PKR activation and eIF2α signaling leading to a compromised stress response, Neurobiology of Disease 146: 105135.

Burnett, S. B., Vaughn, L. S., Strom, J. M., Francois, A. and Patel, R. C. (2019) A truncated PACT protein resulting from a frameshift mutation reported in movement disorder DYT16 triggers caspase activation and apoptosis, Journal of Cellular Biochemistry 120(11): 19004–19018.

Burwick, N., Fang, X., Chorev, M. and Aktas, B. H. (2026) The heme-regulated inhibitor eIF2α kinase: a multifaceted sensor and drug target, Trends in Cell Biology.

Caffall, Z. F., Wilkes, B. J., Hernández-Martinez, R., Rittiner, J. E., Fox, J. T., Wan, K. K., Shipman, M. K., Titus, S. A., Zhang, Y. Q., Patnaik, S. et al. (2021) The HIV protease inhibitor, ritonavir, corrects diverse brain phenotypes across development in mouse model of DYT-TOR1A dystonia, Science Translational Medicine 13(607).

Calakos, N. and Caffall, Z. F. (2024) The integrated stress response pathway and neuromodulator signaling in the brain: lessons learned from dystonia, Journal of Clinical Investigation 134(7).

Calakos, N. and Zech, M. (2025) Emerging Molecular-Genetic Families in Dystonia: Endosome-Autophagosome-Lysosome and Integrated Stress Response Pathways, Movement Disorders 40(1): 7–21.

Camargos, S., Lees, A. J., Singleton, A. and Cardoso, F. (2012) DYT16: the original cases, J Neurol Neurosurg Psychiatry 83(10): 1012–1014.

Camargos, S., Scholz, S., Simon-Sanchez, J., Paisan-Ruiz, C., Lewis, P., Hernandez, D., Ding, J., Gibbs, J. R., Cookson, M. R., Bras, J. et al. (2008) DYT16, a novel young-onset dystonia-parkinsonism disorder: identification of a segregating mutation in the stress-response protein PRKRA, Lancet Neurology 7(3): 207–215.

Caporali, S., De Stefano, A., Calabrese, C., Giovannelli, A., Pieri, M., Savini, I., Tesauro, M., Bernardini, S., Minieri, M. and Terrinoni, A. (2022) Anti-Inflammatory and Active Biological Properties of the Plant-Derived Bioactive Compounds Luteolin and Luteolin 7-Glucoside, Nutrients 14(6).

Cesaro, T. and Michiels, T. (2021) Inhibition of PKR by Viruses, Frontiers in Microbiology 12: 757238.

Chen, D., Fan, Z., Rauh, M., Buchfelder, M., Eyupoglu, I. Y. and Savaskan, N. (2017) ATF4 promotes angiogenesis and neuronal cell death and confers ferroptosis in a xCT-dependent manner, Oncogene 36(40): 5593–5608.

Chen, H., Liu, D., Hua, J., Wu, M., Hua, Y., Feng, C., He, Z., Moffett, P. and Zhang, K. (2026) Hijacking the unfolded protein response (UPR) pathway: Balancing viral infection and host cell survival, Journal of Biological Chemistry 302(1): 111040.

Chen, H. M., Wang, L. and D’Mello, S. R. (2008) A chemical compound commonly used to inhibit PKR, {8- (imidazol-4-ylmethylene)-6H-azolidino[5,4-g] benzothiazol-7-one}, protects neurons by inhibiting cyclin-dependent kinase, European Journal of Neuroscience 28(10): 2003–2016.

Chukwurah, E., Farabaugh, K. T., Guan, B. J., Ramakrishnan, P. and Hatzoglou, M. (2021) A tale of two proteins: PACT and PKR and their roles in inflammation, FEBS Journal.

Clemens, M. J. and Elia, A. (1997) The double-stranded RNA-dependent protein kinase PKR: structure and function, J Interferon Cytokine Res 17(9): 503–524.

Cole, J. L. (2007) Activation of PKR: an open and shut case?, Trends in Biochemical Sciences 32(2): 57–62.

Costa-Mattioli, M. and Walter, P. (2020) The integrated stress response: From mechanism to disease, Science 368(6489).

Dabo, S. and Meurs, E. F. (2012) dsRNA-dependent protein kinase PKR and its role in stress, signaling and HCV infection, Viruses 4(11): 2598–2635.

Dauber, B. and Wolff, T. (2009) Activation of the Antiviral Kinase PKR and Viral Countermeasures, Viruses 1(3): 523–544.

de Carvalho Aguiar, P., Borges, V., Ferraz, H. B. and Ozelius, L. J. (2015) Novel compound heterozygous mutations in PRKRA cause pure dystonia, Movement Disorders 30(6): 877–878.

Demmings, M. D., Kane, E. A., Tennyson, E. C., Hurley, K., Zhao, J., Cruickshanks, N. A., Ciz, V., Krupa, J. M., Pasternak, S. H. and Cregan, S. P. (2026) ATF4 activates a transcriptional program that chronically suppresses mTOR activity and promotes neurodegeneration in Parkinson disease models, Cell Reports 45(8): 117832.

Demmings, M. D., Tennyson, E. C., Petroff, G. N., Tarnowski-Garner, H. E. and Cregan, S. P. (2021) Activating transcription factor-4 promotes neuronal death induced by Parkinson’s disease neurotoxins and α-synuclein aggregates, Cell Death Differ 28(5): 1627–1643.

Dos Santos, C. O., da Silva-Junior, F. P., Puga, R. D., Barbosa, E. R., Azevedo Silva, S. M. C., Borges, V., Limongi, J. C. P., Rocha, M. S. G., Ferraz, H. B. and de Carvalho Aguiar, P. (2018) The prevalence of PRKRA mutations in idiopathic dystonia, Parkinsonism & Related Disorders 48: 93–96.

Farabaugh, K. T., Krokowski, D., Guan, B. J., Gao, Z., Gao, X. H., Wu, J., Jobava, R., Ray, G., de Jesus, T. J., Bianchi, M. G. et al. (2020) PACT-mediated PKR activation acts as a hyperosmotic stress intensity sensor weakening osmoadaptation and enhancing inflammation, Elife 9.

Farabaugh, K. T., Majumder, M., Guan, B. J., Jobava, R., Wu, J., Krokowski, D., Gao, X. H., Schuster, A., Longworth, M., Chan, E. D. et al. (2017) Protein Kinase R Mediates the Inflammatory Response Induced by Hyperosmotic Stress, Mol Cell Biol 37(4).

Frederick, K. and Patel, R. C. (2023) Luteolin protects DYT-PRKRA cells from apoptosis by suppressing PKR activation, Frontiers in Pharmacology 14: 1118725.

Galehdar, Z., Swan, P., Fuerth, B., Callaghan, S. M., Park, D. S. and Cregan, S. P. (2010) Neuronal apoptosis induced by endoplasmic reticulum stress is regulated by ATF4-CHOP-mediated induction of the Bcl-2 homology 3-only member PUMA, Journal of Neuroscience 30(50): 16938–16948.

Garcia, M. A., Gil, J., Ventoso, I., Guerra, S., Domingo, E., Rivas, C. and Esteban, M. (2006) Impact of protein kinase PKR in cell biology: from antiviral to antiproliferative action, Microbiol Mol Biol Rev 70(4): 1032–1060.

Gardner, B. M., Pincus, D., Gotthardt, K., Gallagher, C. M. and Walter, P. (2013) Endoplasmic reticulum stress sensing in the unfolded protein response, Cold Spring Harbor Perspectives in Biology 5(3): a013169.

Gomes, Ilds, Amorim, E. D., León, J. S. S., Schlickmann, T. H., Borges, Y. F. F., Schuh, A. F. S., Teixeira-Dos-Santos, D., Trentin, S., Monte, T. L. and de Oliveira, B. S. F. (2026) Adult-Onset Dystonia Related to PRKRA (DYT16): A Predictor of Favorable Deep Brain Stimulation Outcome, Mov Disord Clin Pract.

Gonzalez-Latapi, P., Marotta, N. and Mencacci, N. E. (2021) Emerging and converging molecular mechanisms in dystonia, J Neural Transm (Vienna) 128(4): 483–498.

Heifetz, A., Keenan, R. W. and Elbein, A. D. (1979) Mechanism of action of tunicamycin on the UDP-GlcNAc:dolichyl-phosphate Glc-NAc-1-phosphate transferase, Biochemistry 18(11): 2186–2192.

Helseth, A. R., Hernandez-Martinez, R., Hall, V. L., Oliver, M. L., Turner, B. D., Caffall, Z. F., Rittiner, J. E., Shipman, M. K., King, C. S., Gradinaru, V. et al. (2021) Cholinergic neurons constitutively engage the ISR for dopamine modulation and skill learning in mice, Science 372(6540).

Hu, Y. and Conway, T. W. (1993) 2-Aminopurine inhibits the double-stranded RNA-dependent protein kinase both in vitro and in vivo, Journal of Interferon Research 13(5): 323–328.

Huang, S., Qu, L. K., Cuddihy, A. R., Ragheb, R., Taya, Y. and Koromilas, A. E. (2003) Protein kinase inhibitor 2-aminopurine overrides multiple genotoxic stress-induced cellular pathways to promote cell survival, Oncogene 22(24): 3721–3733.

Huang, X., Hutchins, B. and Patel, R. C. (2002) The C-terminal, third conserved motif of the protein activator PACT plays an essential role in the activation of double-stranded-RNA-dependent protein kinase (PKR), Biochemical Journal 366(Pt 1): 175–186.

Ingrand, S., Barrier, L., Lafay-Chebassier, C., Fauconneau, B., Page, G. and Hugon, J. (2007) The oxindole/imidazole derivative C16 reduces in vivo brain PKR activation, FEBS Letters 581(23): 4473–4478.

Ito, T., Yang, M. and May, W. S. (1999) RAX, a cellular activator for double-stranded RNA-dependent protein kinase during stress signaling, Journal of Biological Chemistry 274(22): 15427–15432.

Jinnah, H. A. and Sun, Y. V. (2019) Dystonia genes and their biological pathways, Neurobiology of Disease 129: 159–168.

Kojima, E., Takeuchi, A., Haneda, M., Yagi, A., Hasegawa, T., Yamaki, K., Takeda, K., Akira, S., Shimokata, K. and Isobe, K. (2003) The function of GADD34 is a recovery from a shutoff of protein synthesis induced by ER stress: elucidation by GADD34-deficient mice, FASEB Journal 17(11): 1573–1575.

Kok, K. H., Lui, P. Y., Ng, M. H., Siu, K. L., Au, S. W. and Jin, D. Y. (2011) The double-stranded RNA-binding protein PACT functions as a cellular activator of RIG-I to facilitate innate antiviral response, Cell Host & Microbe 9(4): 299–309.

Kolbel, H., Kuechler, A., Strom, T. M., Ludecke, H. J., Moller-Hartmann, C., Della-Marina, A., Wieczoreck, D. and Schara, U. (2017) DYT16 mimics metabolic disease with fever associated beginning of dystonia and MRI abnormalities., European Journal of Pediatric Neurology 21: e177.

Korennykh, A. and Walter, P. (2012) Structural basis of the unfolded protein response, Annu Rev Cell Dev Biol 28: 251–277.

Kuipers, D. J. S., Mandemakers, W., Lu, C. S., Olgiati, S., Breedveld, G. J., Fevga, C., Tadic, V., Carecchio, M., Osterman, B., Sagi-Dain, L. et al. (2021) EIF2AK2 Missense Variants Associated with Early Onset Generalized Dystonia, Annals of Neurology 89(3): 485–497.

Lange, L. M., Junker, J., Loens, S., Baumann, H., Olschewski, L., Schaake, S., Madoev, H., Petkovic, S., Kuhnke, N., Kasten, M. et al. (2021) Genotype-Phenotype Relations for Isolated Dystonia Genes: MDSGene Systematic Review, Movement Disorders 36(5): 1086–1103.

Lee, E. S., Yoon, C. H., Kim, Y. S. and Bae, Y. S. (2007) The double-strand RNA-dependent protein kinase PKR plays a significant role in a sustained ER stress-induced apoptosis, FEBS Letters 581(22): 4325–4332.

Lemmon, M. E., Lavenstein, B., Applegate, C. D., Hamosh, A., Tekes, A. and Singer, H. S. (2013) A novel presentation of DYT 16: acute onset in infancy and association with MRI abnormalities, Movement Disorders 28(14): 1937–1938.

Li, Y., Guo, Y., Tang, J., Jiang, J. and Chen, Z. (2014) New insights into the roles of CHOP-induced apoptosis in ER stress, Acta Biochim Biophys Sin (Shanghai) 46(8): 629–640.

Lockshin, E. R. and Calakos, N. (2024) The integrated stress response in brain diseases: A double-edged sword for proteostasis and synapses, Current Opinion in Neurobiology 87: 102886.

Lui, P. Y., Wong, L. R., Ho, T. H., Au, S. W. N., Chan, C. P., Kok, K. H. and Jin, D. Y. (2017) PACT Facilitates RNA-Induced Activation of MDA5 by Promoting MDA5 Oligomerization, Journal of Immunology 199(5): 1846–1855.

Lytton, J., Westlin, M. and Hanley, M. R. (1991) Thapsigargin inhibits the sarcoplasmic or endoplasmic reticulum Ca-ATPase family of calcium pumps, Journal of Biological Chemistry 266(26): 17067–17071.

Ma, Y. and Hendershot, L. M. (2003) Delineation of a negative feedback regulatory loop that controls protein translation during endoplasmic reticulum stress, Journal of Biological Chemistry 278(37): 34864–34873.

Mao, D., Reuter, C. M., Ruzhnikov, M. R. Z., Beck, A. E., Farrow, E. G., Emrick, L. T., Rosenfeld, J. A., Mackenzie, K. M., Robak, L., Wheeler, M. T. et al. (2020) De novo EIF2AK1 and EIF2AK2 Variants Are Associated with Developmental Delay, Leukoencephalopathy, and Neurologic Decompensation, American Journal of Human Genetics 106(4): 570–583.

Marchal, J. A., Lopez, G. J., Peran, M., Comino, A., Delgado, J. R., Garcia-Garcia, J. A., Conde, V., Aranda, F. M., Rivas, C., Esteban, M. et al. (2014) The impact of PKR activation: from neurodegeneration to cancer, FASEB Journal 28(5): 1965–1974.

Marciniak, S. J., Garcia-Bonilla, L., Hu, J., Harding, H. P. and Ron, D. (2006) Activation-dependent substrate recruitment by the eukaryotic translation initiation factor 2 kinase PERK, Journal of Cell Biology 172(2): 201–209.

Marciniak, S. J., Yun, C. Y., Oyadomari, S., Novoa, I., Zhang, Y., Jungreis, R., Nagata, K., Harding, H. P. and Ron, D. (2004) CHOP induces death by promoting protein synthesis and oxidation in the stressed endoplasmic reticulum, Genes Dev 18(24): 3066–3077.

Masnada, S., Martinelli, D., Correa-Vela, M., Agolini, E., Baide-Mairena, H., Marcé-Grau, A., Parazzini, C., Veggiotti, P., Perez-Duenas, B. and Tonduti, D. (2021) PRKRA-Related Disorders: Bilateral Striatal Degeneration in Addition to DYT16 Spectrum, Movement Disorders 36(4): 1038–1040.

Najmabadi, H., Hu, H., Garshasbi, M., Zemojtel, T., Abedini, S. S., Chen, W., Hosseini, M., Behjati, F., Haas, S., Jamali, P. et al. (2011) Deep sequencing reveals 50 novel genes for recessive cognitive disorders, Nature 478(7367): 57–63.

Neill, G. and Masson, G. R. (2023) A stay of execution: ATF4 regulation and potential outcomes for the integrated stress response, Frontiers in Molecular Neuroscience 16: 1112253.

Oyadomari, S., Koizumi, A., Takeda, K., Gotoh, T., Akira, S., Araki, E. and Mori, M. (2002) Targeted disruption of the Chop gene delays endoplasmic reticulum stress-mediated diabetes, Journal of Clinical Investigation 109(4): 525–532.

Pakos-Zebrucka, K., Koryga, I., Mnich, K., Ljujic, M., Samali, A. and Gorman, A. M. (2016) The integrated stress response, EMBO Rep 17(10): 1374–1395.

Patel, C. V., Handy, I., Goldsmith, T. and Patel, R. C. (2000) PACT, a stress-modulated cellular activator of interferon-induced double-stranded RNA-activated protein kinase, PKR, Journal of Biological Chemistry 275(48): 37993–37998.

Patel, R. C. and Sen, G. C. (1998) PACT, a protein activator of the interferon-induced protein kinase, PKR, EMBO Journal 17(15): 4379–4390.

Pinto, M. J., Oliveira, A., Rosas, M. J. and Massano, J. (2020) Imaging Evidence of Nigrostriatal Degeneration in DYT-PRKRA, Mov Disord Clin Pract 7(4): 472–474.

Pitale, P. M., Gorbatyuk, O. and Gorbatyuk, M. (2017) Neurodegeneration: Keeping ATF4 on a Tight Leash, Frontiers in Cellular Neuroscience 11: 410.

Quadri, M., Olgiati, S., Sensi, M., Gualandi, F., Groppo, E., Rispoli, V., Graafland, J., Breedveld, G. J., Fabbrini, G., Berardelli, A. et al. (2016) PRKRA Mutation Causing Early-Onset Generalized Dystonia-Parkinsonism (DYT16) in an Italian Family, Movement Disorders 31(5): 765–767.

Rittiner, J. E., Caffall, Z. F., Hernandez-Martinez, R., Sanderson, S. M., Pearson, J. L., Tsukayama, K. K., Liu, A. Y., Xiao, C., Tracy, S., Shipman, M. K. et al. (2016) Functional Genomic Analyses of Mendelian and Sporadic Disease Identify Impaired eIF2alpha Signaling as a Generalizable Mechanism for Dystonia, Neuron 92(6): 1238–1251.

Samuel, C. E., Kuhen, K. L., George, C. X., Ortega, L. G., Rende-Fournier, R. and Tanaka, H. (1997) The PKR protein kinase--an interferon-inducible regulator of cell growth and differentiation, International Journal of Hematology 65(3): 227–237.

Sano, R. and Reed, J. C. (2013) ER stress-induced cell death mechanisms, Biochim Biophys Acta 1833(12): 3460–3470.

Schroder, M. and Kaufman, R. J. (2005) The mammalian unfolded protein response, Annual Review of Biochemistry 74: 739–789.

Seibler, P., Djarmati, A., Langpap, B., Hagenah, J., Schmidt, A., Bruggemann, N., Siebner, H., Jabusch, H. C., Altenmuller, E., Munchau, A. et al. (2008) A heterozygous frameshift mutation in PRKRA (DYT16) associated with generalised dystonia in a German patient, Lancet Neurology 7(5): 380–381.

Simon, Tricia A. and Patel, Rekha C. (2025) Molecular mechanisms in DYT-PRKRA: pathways regulated by PKR activator protein PACT, Dystonia Volume 4–2025.

Singh, M., Castillo, D., Patel, C. V. and Patel, R. C. (2011) Stress-induced phosphorylation of PACT reduces its interaction with TRBP and leads to PKR activation, Biochemistry 50(21): 4550–4560.

Singh, M., Fowlkes, V., Handy, I., Patel, C. V. and Patel, R. C. (2009) Essential role of PACT-mediated PKR activation in tunicamycin-induced apoptosis, Journal of Molecular Biology 385(2): 457–468.

Singh, M. and Patel, R. C. (2012) Increased interaction between PACT molecules in response to stress signals is required for PKR activation, Journal of Cellular Biochemistry 113(8): 2754–2764.

Soleymani, F., Piryaei, F., Farhadi, A., Pourbakhtyaran, E. and Behroozi, J. (2025) A Homozygous c.74A>G Variant in PRKRA Causes DYT-PRKRA: Extensive Familial Segregation and a Variant of Uncertain Significance (VUS) Reclassification, Movement Disorders 40(10): 2257–2261.

Sun, X., Liu, J., Crary, J. F., Malagelada, C., Sulzer, D., Greene, L. A. and Levy, O. A. (2013) ATF4 protects against neuronal death in cellular Parkinson’s disease models by maintaining levels of parkin, Journal of Neuroscience 33(6): 2398–2407.

Szaruga, M., Janssen, D. A., de Miguel, C., Hodgson, G., Fatalska, A., Pitera, A. P., Andreeva, A. and Bertolotti, A. (2023) Activation of the integrated stress response by inhibitors of its kinases, Nat Commun 14(1): 5535.

Thomsen, M., Lange, L. M., Zech, M. and Lohmann, K. (2024) Genetics and Pathogenesis of Dystonia, Annual Review of Pathology 19: 99–131.

Tisch, S. and Kumar, K. R. (2020) Pallidal Deep Brain Stimulation for Monogenic Dystonia: The Effect of Gene on Outcome, Frontiers in Neurology 11: 630391.

Tronel, C., Page, G., Bodard, S., Chalon, S. and Antier, D. (2014) The specific PKR inhibitor C16 prevents apoptosis and IL-1beta production in an acute excitotoxic rat model with a neuroinflammatory component, Neurochemistry International 64: 73–83.

Vaughn, L. S., Bragg, D. C., Sharma, N., Camargos, S., Cardoso, F. and Patel, R. C. (2015) Altered Activation of Protein Kinase PKR and Enhanced Apoptosis in Dystonia Cells Carrying a Mutation in PKR Activator Protein PACT, Journal of Biological Chemistry 290(37): 22543–22557.

Vaughn, L. S., Frederick, K., Burnett, S. B., Sharma, N., Bragg, D. C., Camargos, S., Cardoso, F. and Patel, R. C. (2022) DYT-PRKRA Mutation P222L Enhances PACT’s Stimulatory Activity on Type I Interferon Induction, Biomolecules 12(5).

Volchuk, A. and Ron, D. (2010) The endoplasmic reticulum stress response in the pancreatic β-cell, Diabetes, Obesity & Metabolism 12 Suppl 2: 48–57.

Walter, P. and Ron, D. (2011) The unfolded protein response: from stress pathway to homeostatic regulation, Science 334(6059): 1081–1086.

Wang, X., Zhao, J., Li, Y., Gao, S. and Su, L. (2025) Luteolin as a multifaceted immunomodulator: insights into its effects on diverse immune cell populations and therapeutic implications, Frontiers in Immunology 16: 1621367.

Wek, R. C., Jiang, H. Y. and Anthony, T. G. (2006) Coping with stress: eIF2 kinases and translational control, Biochemical Society Transactions 34(Pt 1): 7–11.

Wortel, I. M. N., van der Meer, L. T., Kilberg, M. S. and van Leeuwen, F. N. (2017) Surviving Stress: Modulation of ATF4-Mediated Stress Responses in Normal and Malignant Cells, Trends Endocrinol Metab 28(11): 794–806.

Xiao, J., Vemula, S. R., Xue, Y., Khan, M. M., Carlisle, F. A., Waite, A. J., Blake, D. J., Dragatsis, I., Zhao, Y. and LeDoux, M. S. (2017) Role of major and brain-specific Sgce isoforms in the pathogenesis of myoclonus-dystonia syndrome, Neurobiology of Disease 98: 52–65.

Yan, W., Frank, C. L., Korth, M. J., Sopher, B. L., Novoa, I., Ron, D. and Katze, M. G. (2002) Control of PERK eIF2alpha kinase activity by the endoplasmic reticulum stress-induced molecular chaperone P58IPK, Proceedings of the National Academy of Sciences of the United States of America 99(25): 15920–15925.

Young, S. K. and Wek, R. C. (2016) Upstream Open Reading Frames Differentially Regulate Gene-specific Translation in the Integrated Stress Response, Journal of Biological Chemistry 291(33): 16927–16935.

Zakirova, Z., Fanutza, T., Bonet, J., Readhead, B., Zhang, W., Yi, Z., Beauvais, G., Zwaka, T. P., Ozelius, L. J., Blitzer, R. D. et al. (2018) Mutations in THAP1/DYT6 reveal that diverse dystonia genes disrupt similar neuronal pathways and functions, PLoS Genetics 14(1): e1007169.

Zech, M., Castrop, F., Schormair, B., Jochim, A., Wieland, T., Gross, N., Lichtner, P., Peters, A., Gieger, C., Meitinger, T. et al. (2014) DYT16 revisited: exome sequencing identifies PRKRA mutations in a European dystonia family, Movement Disorders 29(12): 1504–1510.

Zhang, L. and Wang, A. (2012) Virus-induced ER stress and the unfolded protein response, Front Plant Sci 3: 293.

Zhu, J. and Marciniak, S. J. (2026) GCN2 in proteostasis: structural logic, signalling networks and disease, FEBS Journal 293(18): 5627–5644.

